# Skeletal muscle MACF1 maintains myonuclei and mitochondria localization through microtubules to control muscle functionalities

**DOI:** 10.1101/636464

**Authors:** Alireza Ghasemizadeh, Emilie Christin, Alexandre Guiraud, Nathalie Couturier, Valérie Risson, Emmanuelle Girard, Christophe Jagla, Cedric Soler, Lilia Laddada, Colline Sanchez, Francisco Jaque, Audrey Garcia, Marine Lanfranchi, Vincent Jacquemond, Julien Gondin, Julien Courchet, Laurent Schaeffer, Vincent Gache

## Abstract

Skeletal muscle is made from multinuclear myofiber, where myonuclei are positioned at the periphery or clustered below neuromuscular junctions (NMJs). While mispositioned myonuclei are the hallmark of numerous muscular diseases, the molecular machinery maintaining myonuclei positioning in mature muscle is still unknown. Here, we identified microtubule-associated protein MACF1 as an evolutionary conserved regulator of myonuclei positioning, in vitro and in vivo, controlling the “microtubule code” and stabilizing the microtubule dynamics during myofibers maturation, preferentially at NMJs. Specifically, MACF1 governs myonuclei motion, mitochondria positioning and structure and acetylcholine receptors (AChRs) clustering. Macf1-KO in young and adult mice decreases muscle excitability and causes evolutionary myonuclei positioning alterations in adult mice, paralleled with high mitochondria content and improved resistance to fatigue. We present MACF1 as a primary actor of the maintenance of synaptic myonuclei and AChRs clustering, peripheral myonuclei positioning and mitochondria organization through the control of microtubule network dynamics in muscle fibers.

## Introduction

Throughout muscle development, myonuclei actively position themselves and adapt special localization in mature myofibers where myonuclei are regularly spaced along muscle fibers and localized at the periphery (Roman and Gomes, 2017). The density of myonuclei along myofibers’ length remains relatively constant with the exception of immediately below the neuromuscular junction (NMJ) where about 5-6 myonuclei (synaptic myonuclei) are clustered (Manhart et al., 2018; Ravel-Chapuis et al., 2007; Bruusgaard et al., 2006). Failure in extra-synaptic myonuclei patterning has been linked to different myopathies and muscle weakness (Metzger et al., 2012; Roman et al., 2017; Falcone et al., 2014; Collins et al., 2017; Perillo and Folker, 2018). This precise organization of myonuclei is correlated with particular myonuclei shape in muscle fibers, the alteration of which has recently emerged as potentially contributing to several muscular diseases (Norton and Phillips-Cremins, 2017; Cho et al., 2017; Puckelwartz et al., 2009; Méjat et al., 2009; Janin and Gache, 2018).

Proteins localized inside myonuclei such as Lamin A/C, Emerin or Nuclear Envelope Proteins (NEPs), that couple the nuclear lamina to cytoskeleton through the “LInker of Nucleoskeleton and Cytoskeleton” (LINC) complex, play critical roles in the maintenance of myonuclei shape and localization (Chang et al., 2015; Lee and Burke, 2017; Levy et al., 2018; Mattioli et al., 2011). Additionally, molecular motors, Microtubule-Associated Proteins (MAPs), in interaction with actin/microtubule networks are also involved in myonuclei localization during muscle formation (Gache et al., 2017; Rivero et al., 2009; Casey et al., 2003; Metzger et al., 2012). Most importantly, this precise localization and shape, determined by mechanical forces emanating from different cytoskeletons networks, acts directly on myonuclei-triggered chromatin organization and gene expression (Ramdas and Shivashankar, 2015; Robson et al., 2016; Chojnowski et al., 2015; Kirby and Lammerding, 2018). Still, how late positioning of myonuclei in myofibers is set and how it regulates signaling within cells that contributes to different pathways to maintain muscle integrity are still poorly understood.

To identify proteins involved in the late positioning of myonuclei, we used primary myotubes/myofibers culture and identified MACF1 (Microtubule Actin Cross Factor 1) as a regulator of myonuclei positioning in late phases of myofiber maturation. MACF1 is a member of the plakin family. It is a large protein composed of multiple domains capable of interacting with actin filaments and microtubules that helps stabilizing them towards adhesion structures in a variety of tissues (Kodama et al., 2003; Sanchez-Soriano et al., 2009). In addition, other supporting evidences have shown a role of MACF1 in myonuclei position. The *C. elegans* orthologous, VAB-10 and the *Drosophila* orthologous, Shot, are essential for nuclei migration in distal tip cells of the somatic gonad and myotendinous junction formation respectively (Kim et al., 2011; Lee and Kolodziej, 2002; Bottenberg et al., 2009). Nevertheless, a duplication of *MACF1* gene (inducing a decrease in the total amount of MACF1 protein) has been linked to a new neuromuscular disease (Jørgensen et al., 2014). Also important in this respect, it was recently proposed that Shot/MACF1 participates in the formation and maintenance of a perinuclear shield in the muscle of drosophila larvae, contributing once again in myonuclei localization (Wang et al., 2015).

Here, we demonstrate *in vitro* and *in vivo* that MACF1 is implicated in the maintenance of myonuclei patterning in myofibers, starting at the NMJs, and controlling both myonuclei and mitochondria dynamics through the regulation of microtubules.

## Results

### MACF1 is not required in precocious myonuclei positioning but is essential during myofibers maturation in the maintenance of myonuclei localization

To identify new factors that contribute specifically to myonuclei spreading in myofibers, we purified proteins able to bind to microtubules, stabilized with Taxol^®^, from 3 days mouse primary myotubes (Fig1A). The major Microtubule Associated Protein (MAP) identified by mass-spectrometry using this protocol revealed the significant presence of MACF1/ACF7 protein, a member of the plakin family (Supplementary Fig1A-B). Although MACF1 is initially described to be predominantly expressed in neurons and muscle (Bernier et al., 2000), its role during muscle fibers formation and behavior is poorly understood. Published work suggests that plakin family members play specific roles through the regulation of placement and function of specific organelles such as nucleus, mitochondria, Golgi apparatus, and sarcoplasmic reticulum (Boyer et al., 2010). Mouse *Macf1* mRNA was previously show to increase steadily during myogenesis (Sun et al., 1999) and MACF1 was proposed to contribute in muscle fiber to form a flexible perinuclear shield in collaboration with two other MAPs named EB1 and Nesprin (Wang et al., 2015). We confirmed the increase of *Macf1* mRNA and MACF1 protein during early steps of myotubes formation (3 to 5 days of differentiation) using RT-qPCR and Western blotting on mouse myoblast C2C12 cells (Supplementary Fig1C-D).

**Figure 1:**
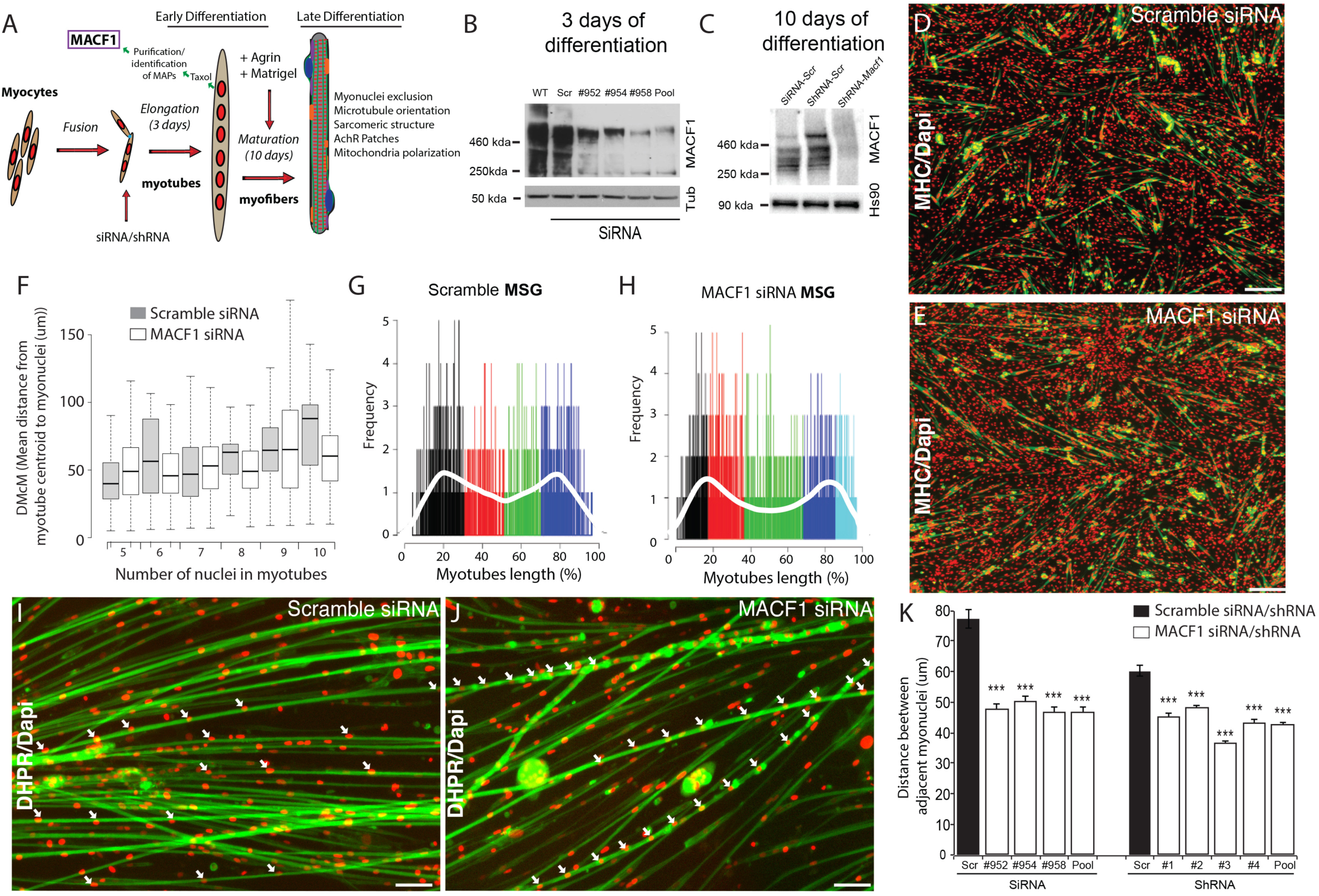
MACF1 is an essential regulator of myonuclei positioning in mature myofibers. (A) Schema of sequential steps in order to obtain immature myotubes and mature myofibers from primary myoblasts. siRNA/shRNA were transfected in early phases of myotubes fusion. (B). Western blot analysis of MACF1 protein expression in total protein extracts of primary myoblast not treated (WT), treated with 3 individual siRNAs (#952, #954, #958) or a pool of 3 individual siRNAs targeting *Macf1* after 3 days of differentiation. Tubulin is used as “loading control”. (C) Western blot analysis of MACF1 protein expression in total protein extracts of primary myoblast treated with a pool of 3 individual siRNAs or a pool of 4 shRNAs targeting *Macf1* after 11 days of differentiation. HSP90 is used as “loading control”. (D-E) Representative images of immunofluorescence staining from 3 individual experiments presenting Myosin Heavy Chain (green, MHC) and myonuclei (red) in primary myotubes treated with scramble (D) or a pool of 3 individual siRNAs targeting *Macf1* (E) after 3 days of differentiation. Scale Bar = 150 µm. (F) Mean distances between each myonuclei and myotube’s centroid ranked by myonuclei content per myotubes were quantified after 3 days of differentiation in cells treated with a scramble or a pool of 3 individual *Macf1* siRNAs. (G-H) Myonuclei Spreading Graph (MSG) represents statistical spatial distribution of myonuclei along myotubes in scramble (G) and in *Macf1* siRNA-treated myotubes (H). In F-H, 3 individual experiments, n=170 for scramble and n=365 for SiRNA *Macf1* conditions. (I-J) Representative images of 10 days differentiated myofibers stained, for DiHydroPyridine Receptor (green) and Dapi (red) in cells treated with scramble siRNA (I) or a pool of 3 individual siRNAs targeting *Macf1* (J). Scale Bar= 50 µm. (K) Mean distance between adjacent myonuclei in primary myotubes treated with scramble, 3 individual siRNAs (#952, #954, #958) or a pool of 3 individual siRNA targeting *Macf11* and in primary myotubes treated with scramble, 4 individual shRNAs (#1, #2, #3, #4) or a pool of 4 individual shRNAs targeting *Macf1* after 10 days of differentiation from 3 individual experiments with at least n=400 by conditions.

We first addressed the role of MACF1 in early steps of myotubes formation (Fig1A). Briefly, isolated murine myoblasts were treated with either short interfering RNA (siRNA) or small hairpin RNA (shRNA) targeting either a scrambled sequence or the *Macf1* gene to obtain a strong reduction of MACF1 expression after 3 or 10 days of differentiation (Fig1B-C). Primary myoblasts were then induced *in vitro* to form myotubes that were fixed and analyzed after 3 days of differentiation (Fig1D-E). At this step, myoblasts have fused together and formed immature myofibers (myotubes) characterized by myonuclei alignment in the middle of myotubes, mainly organized by an interplay between MAP7, Kif5b and microtubules (Metzger et al., 2012). Surprisingly at this early step of differentiation, MACF1 down-regulation had no impact on parameters related to myotubes architecture, reflected by myotubes areas and/or length according to myonuclei content (Supplementary Fig1E-F). Additionally, myonuclei repartition reflected by the mean distance of each myonuclei from myotubes centroid or by the statistical repartition of myonuclei along the length of myotubes, was not affected by the down regulation of MACF1 (Fig1F-H). These results were confirmed in C2C12 myoblasts cells, treated with either siRNA or shRNA targeting *Macf1* and induced to form myotubes (Supplementary Fig1G-J). This suggested that for a defined myonuclei content in myotubes, MACF1 does not act as an early “anti-elongation” factor and is not involved in the process that contributes to the early equilibrium in force spreading emanating on myonuclei in myotubes (Manhart et al., 2018).

To further investigate the implication of MACF1 in late steps of differentiation, primary mice myotubes were maintained in differentiation media for 10 days as previously described (Pimentel et al., 2017; Falcone et al., 2014), covered by matrigel^®^ and supplemented with neural Agrin (Fig1A). In these conditions of maturation, myonuclei are compressed between the membrane of the myofiber and sarcomere structure, adopt a flatten architecture and spread all along myofibers (Roman et al., 2017). This long-term differentiation approach allowed the identification of MACF1 as a regulator of myonuclear positioning in the maturation steps of developing myofibers (Fig1I-J). Actually, MACF1 depletion caused a 40% decrease of the mean distance between adjacent myonuclei using either each of 3 individual siRNA or a pool of individual siRNAs (Fig1K). We confirmed this data with an alternative approach using 4 individual shRNA or a pool of these 4 shRNAs (Fig1K) and observed a 30% decrease in the mean distance between myonuclei.

Altogether, these results suggests that MACF1 is progressively expressed and accumulates during early steps of differentiation processes without implication in the precocious microtubules-dependent localization of myonuclei, yet during myofibers maturation steps, MACF1 gets involved in the maintenance of the distance between adjacent myonuclei.

### MACF1 controls Myonuclei shape, microtubule network, mitochondria localization and AChRs clustering in mature myofibers

MACF1 is a member of the plakin family (Gong et al., 2001; Goryunov and Liem, 2016), composed of multiple domains interacting with and stabilizing both actin filaments and microtubules network (Supplementary Fig1B) in variety of tissues (Kodama et al., 2003; Sanchez-Soriano et al., 2009). To further investigate the specific roles of MACF1 in mature myofibers, we analyzed the structure of actin and microtubule cytoskeleton networks (Fig2A). The structure of sarcomeres reflected by actin and alpha-actinin staining did not seem to be affected in the absence of MACF1, as evidenced by the proper peripheral localization of myonuclei (Fig2A)(Roman et al., 2017). However, the microtubule network was slightly altered all along myofibers with a net architectural modification at the vicinity of myonuclei (Fig2A). To discard an effect of the absence of MACF1 on the global shape of myofibers, we analyzed the width of each individual myofiber and did not observe significant differences between scrambled and *macf1* si/shRNA treated myofibers (Fig2B).

**Figure 2:**
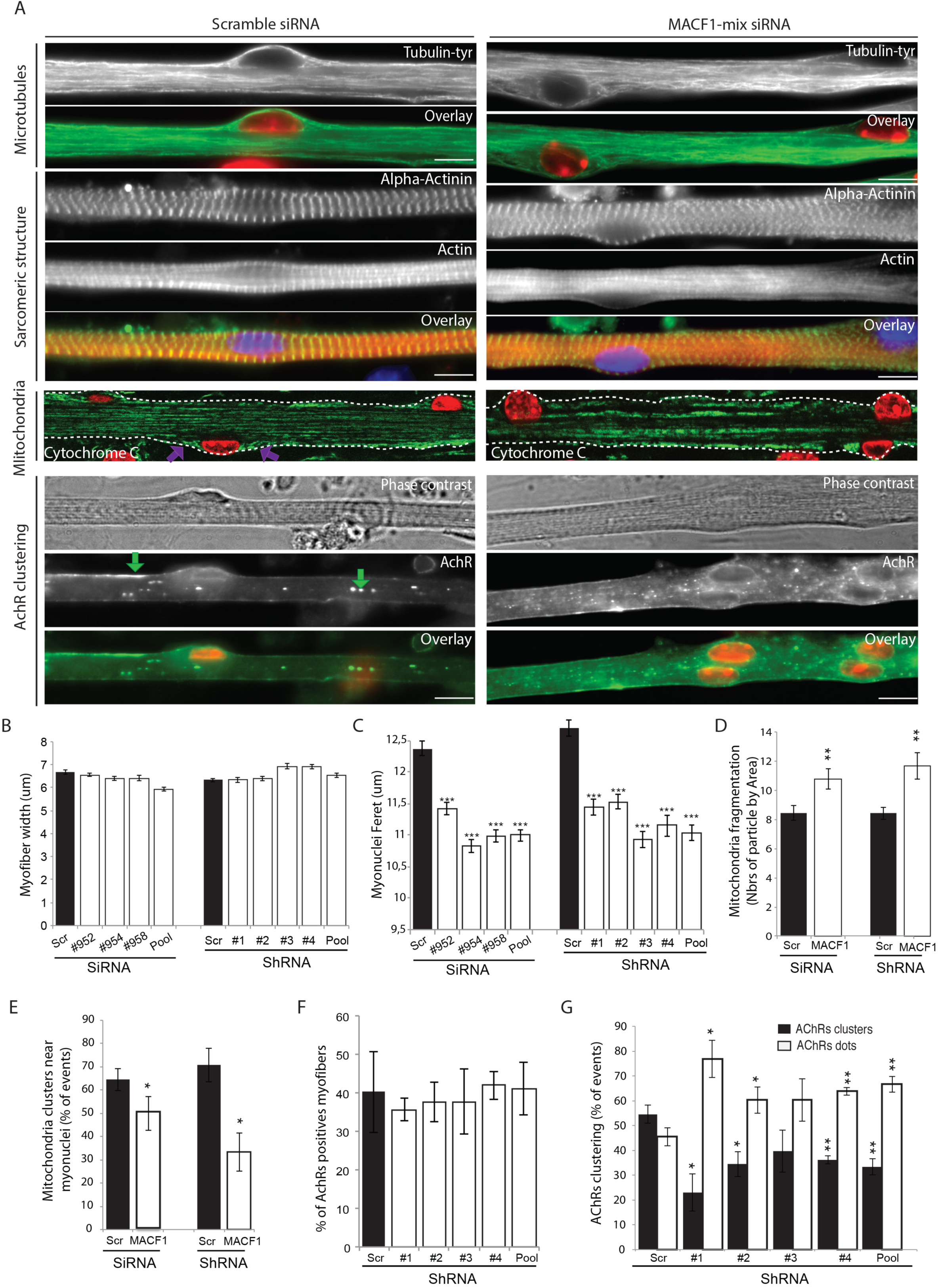
MACF1 control myonuclei shape, microtubule integrity, mitochondria localization and AChRs clustering in mature myofibers. (A) Representative images of 10 days differentiated myofibers, stained, from the top to the bottom respectively, for Tyrosinated-Tubulin (green), Alpha-actinin (green), Actin (red), Cytochrome-C (green) and acetylcholine receptor (Green) in cells treated with scramble siRNA (left panels) or a pool of 3 individual siRNAs targeting *Macf1* (right panel). Myonuclei are stained in blue or red. Scale Bar= 20 µm. (B) Myofibers width (3 individual experiments with at least n=190 by conditions) and (C) myonuclei ferret (3 individual experiments with at least n=170 by conditions) in primary myotubes treated with scramble, 3 individual siRNAs (#952, #954, #958) or a pool of 3 individual siRNAs targeting *Macf1* and in primary myotubes treated with scramble, 4 individual shRNAs (#1, #2, #3, #4) or a pool of 4 individual shRNAs targeting *Macf1* after 10 days of differentiation. (D) Mitochondria fragmentation (2 individual experiments with at least n=800 by conditions) and (E) distribution around myonuclei (2 individual experiments with at least n=30 by conditions) in cells treated with scramble siRNA/shRNA, a pool of 3 individual siRNAs or a pool of 4 individuals shRNAs targeting *Macf1* after 10 days of differentiation. (F) Percentage of myofibers expressing immunofluorescence staining for acetylcholine receptors in primary myotubes treated with scramble shRNA, 4 individual shRNAs (#1, #2, #3, #4) or a pool of 4 individual shRNAs targeting *Macf1* after 10 days of differentiation. 3 individual experiments with at least n=130 by conditions. (G) Acetylcholine receptors (AChR) clustering distribution in cells treated with scramble shRNA or 4 individual shRNAs (#1, #2, #3, #4) or a pool of 4 individual shRNAs targeting *Macf1* after 10 days of differentiation. 3 individual experiments with at least n=130 by condition. Error bars, s.e.m. ***p < 0.001, **p < 0.01, *p < 0.05.

The obvious phenotype regarding myonuclei aspect was an increase of the apparent sphericity of myonuclei (Fig2C). We measured the mean feret diameter of each myonuclei compressed between myofibrils network and the external membrane (Roman et al., 2017) and observed that in mature myofibers, the mean length of myonuclei was decreased by 11 and 16% in the absence of MACF1, using siRNA and shRNA strategies respectively, compared to control conditions (Fig2C).

We also addressed the structure and localization of the mitochondrial network in the presence or absence of MACF1, as revealed by a Cytochrome-C staining in mature myofibers (Fig2A-B). In myofibers treated with scrambled siRNA, mitochondria lined up in the bulk of myofibers, in-between myofibrils, where they followed the microtubule network. However in *Macf1* siRNA/shRNA treated myofibers, we observed a 30% increase of mitochondria fragmentation (Fig2D). Furthermore unlike in control condition where mitochondria clustered in the vicinity of myonuclei with a preference for both extremities of polarized myonuclei (Fig2A, purple arrows), we observed a loss of mitochondria localization at the periphery of myonuclei upon MACF1 knock-down (Fig2E). These results confirmed a role of MACF1 in muscles specifically related to microtubule network alteration, as myonuclei and mitochondria localization were both affected.

Interestingly, the MACF1-dependent microtubule network in charge of both transmitted forces on myonuclei and mitochondria localization, occurred specifically during the maturation process and especially on extracellular matrix treated and Agrin stimulated myofibers. The use of Agrin in this system is known to induce clustering of acetylcholine receptors (AChRs), a synaptic receptor expressed locally at NMJs (Lin et al., 2005; Schmidt et al., 2012; Vilmont et al., 2016) (Fig 2A, green arrows). Thus, we next focused on AChRs clustering and found that even though the proportion of myofibers expressing AChRs was not changed (Fig2F), the ability of myofibers to efficiently cluster AChRs was highly affected resulting in a drop of at least 20% in this clustering ability, determined by the presence of cluster larger than 8 µm (Fig2G). To confirm these results on the potential role of MACF1 on AChRs clustering specifically at the neuromuscular junction, we used the muscle specific driver *(Mef2-GAL4)* to express RNAi against *Drosophila Shot* (*Macf1* orthologous) as described previously (Wang et al., 2015). Immunostaining against Shot confirmed a preferential localization of the protein around myonuclei and showed a significant reduction of protein level around myonuclei in Shot-KD larval muscles (Supplementary Fig2A-B). In accordance with our *in vitro* system, we found that actives zones of synaptic button and post-synaptic terminals were decreased in Shot-KD larval muscles (Supplementary Fig2C-D).

Altogether, these results suggest that MACF1 is implicated in late steps of the myofibers maturation process by maintaining microtubule related spreading of both myonuclei and mitochondria and by controlling AChRs clustering at the NMJs.

### MACF1 controls myonuclei dynamics in a microtubule dependent manner

MACF1 was previously shown to regulate microtubule polymerization (Ka and Kim, 2016; Alves-Silva et al., 2012; Wang et al., 2015). We tested whether in our system microtubule dynamics were changed using the tracking of fluorescently tagged EB1. As expected, EB1 localization at the tips of microtubule was not dependent on the presence of MACF1, however the velocity of EB1 was increased by 20% when MACF1 was down-regulated, confirming the role of MACF1 in the regulation of microtubule polymerization (Fig3A-B; Supplementary movie 1-2). As myonuclei use preferentially the microtubule network to move longitudinally along myotubes (Gache et al., 2017; Metzger et al., 2012), we next investigated the impact of the increase in microtubule dynamicity on myonuclei movements in mature myofibers. Just before fusion occurs, myoblasts were transfected with lamin-chromobody® to allow the visualization of myonuclei concomitantly with shRNA targeting either scramble sequence or *Macf1* with a GFP reporter to select myotubes containing both construction (shRNA and lamin-chromobody®) (Fig3C, Supplementary movie 3-4). In 5-day differentiated myofibers (already covered by matrigel^®^ and supplemented with neural Agrin), myonuclei were tracked every 15 minutes for a time period of 14 hours. Myonuclei displacements parameters were analyzed using SkyPad method (Cadot et al., 2014). In control condition, myonuclei moved during 32 % of the time at a median speed of 0,16 µm/min, resulting in a displacement of 24 µm after 14 hours (more than twice the distance between 2 myonuclei) mainly in the same direction (persistence of 0,98). Interestingly in the absence of MACF1, the median velocity of myonuclei was not significantly changed (Fig3D). On the other hand, the percentage of time that myonuclei spent in motion was doubled to reach 66% of the time (Fig3E). Similarly, the median duration of pause was 530 minutes in the control condition while it fell to 330 minutes in the absence of MACF1 (Fig3F). Interestingly, we also observed a decreased tendency in the persistence of the movement of myonuclei (persistence is 0,68 in absence of MACF1). All in all, in the absence of MACF1, myonuclei travelled twice more distance than in control condition with a mean distance of 44 µm in 14 hours (Fig3G). This result suggests that MACF1, through the increase of microtubule dynamics, increase the movements of myonuclei that will induce a reduction of the distance between adjacent myonuclei.

**Figure 3:**
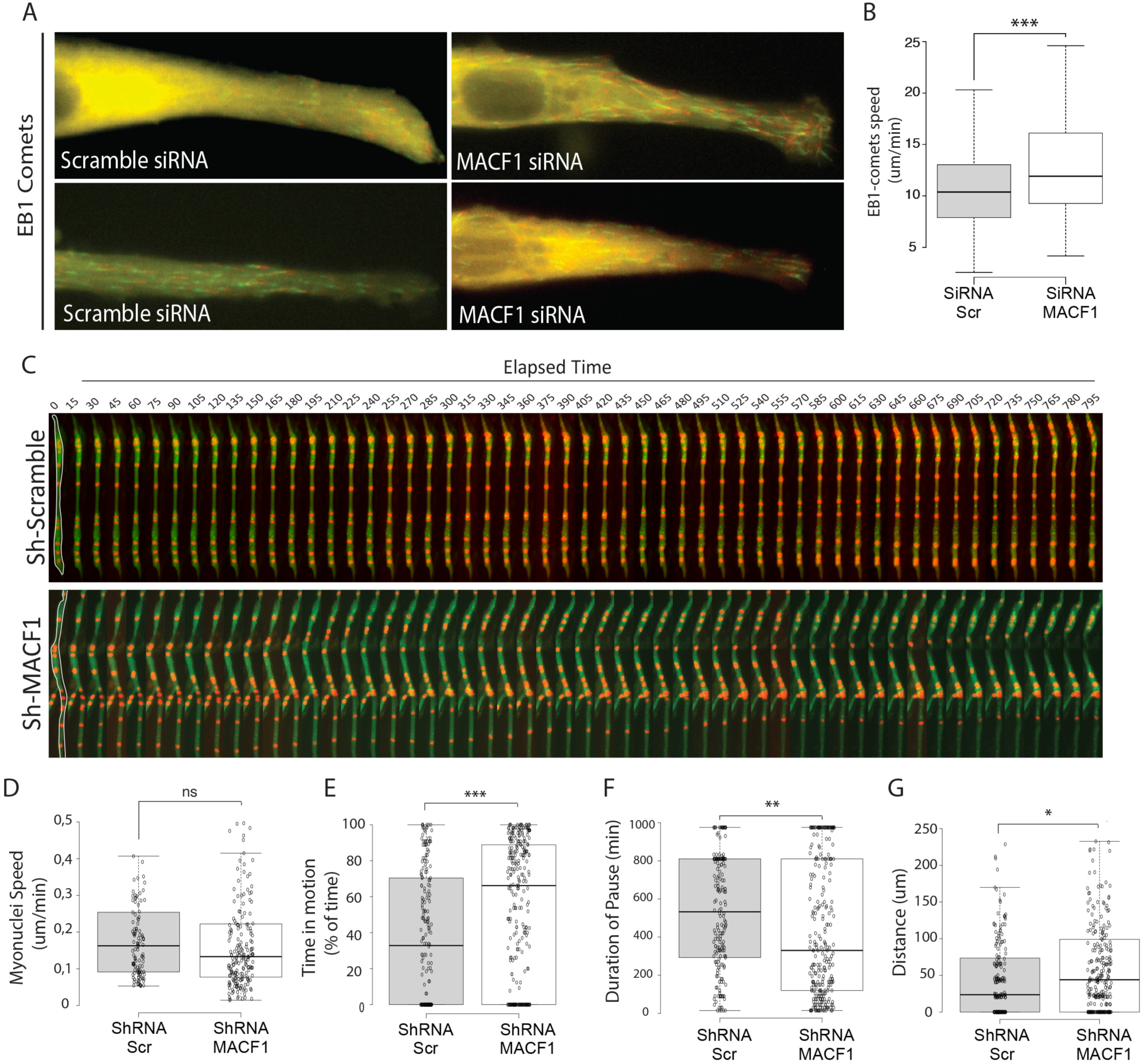
MACF1 controls microtubules and myonuclei dynamics. (A) Representative images of primary myotubes transfected with EB1-GFP with either scramble or a pool of 3 individual siRNAs targeting *Macf1* (#952, #954, #958) showing colored comets. (B) Quantification of EB1 comets speed in myoblasts treated with scramble or a pool of 3 individual siRNAs targeting *Macf1* in primary myotubes after 3 days of differentiation. At least 200 comets were mapped per condition from 3 independent experiments. Error bars, s.e.m. ***p < 0.001. (C) Representatives frames from a 14h time-lapse movie in two channels (shRNA in green and lamin-chromobody® in red) of primary myotubes in the presence of Agrin and covered by matrigel. In the first frame (on the left), myofibers are selected in white, which corresponds to the region used to create the adjacent kymograph. Primary myotubes were treated with scramble or a pool of 4 individual shRNAs targeting *Macf1* after 5 days of differentiation. 3 individual experiments with at least n=200 by conditions. Speed (D), Time in motion (E), Duration of pauses (F) and Distance crossed by myonuclei (G) were quantified using SkyPad analysis (Cadot et al., 2014). ***p < 0.001, **p < 0.01, *p < 0.05.

### MACF1 conditional muscle-KO adult mice exhibit myonuclei mislocalization

To further investigate the role played by MACF1 in skeletal muscles development, we used a conditional *Macf1* knockout mouse line in which exons 6 and 7 of the *Macf1* gene are floxed (Goryunov et al., 2010). Homozygous mice were then crossed with mice carrying the Cre-recombinase expression under the control of the Human Skeletal muscle Actin (HSA)(Miniou et al., 1999). Briefly, mice carrying two floxed alleles of *Macf1* (*Macf1*f/f) were crossed with heterozygous mice for *Macf1* (*Macf1*f/-) that carried Hsa::Cre transgene (Hsa::Cre; *Macf1*f/−), generating the conditional mutant (Hsa::Cre; *Macf1*f/f referred as *Macf1*^f/f^ Cre+) and control mice (WT; *Macf1*f/f referred as *Macf1*^f/f^ Cre-) (Supplementary Fig3A-B). We quantified the loss of *Macf1* by measuring the amount of MACF1 mRNA and protein in muscle tissue lysates (Supplementary Fig3C-D). MACF1 levels were reduced by more than half in muscle tissue, indicating that MACF1 is efficiently removed from myofibers in *Macf1* muscle-conditional mutant mice. We can attribute the remaining presence of MACF1 proteins from other tissues such as fibroblasts, motoneurons and schawnn cells. Mice were born at Mendelian frequencies and appeared indistinguishable from WT littermates (data not shown). The comparative global body weight evolution of conditional mutant (*Macf1*^f/f^ Cre+) and control mice (*Macf1*^f/f^ Cre-) did not show any apparent problems in growing and maturation phase (Supplementary Fig3E).

Myonuclei localization in different type of muscles was studied using a histological staining approach in muscle cross-section. Laminin staining was used to determine the limit of each myofiber and Dapi to detect myonuclei (Fig4A). We observed a dramatic delocalization of myonuclei in 12-month-adult muscle fibers revealed by the presence of myonuclei in the center or dispatched in the myofibers rather than at the periphery in the *Tibialis Anterior, Soleus* and *Gastrocnemius* with a respective increase of 275, 153 and 230% of the delocalization index (Fig4B, Supplementary Fig3F). Interestingly, no significant delocalization of myonuclei was observed in 3- and 8-month-adult mice (Fig4B). At the fiber level, no significant alteration was detected in the distribution of myofibers cross-sectional area (CSA) in 3-months-adult mice in the *Tibialis Anterior* (Fig4C). Nonetheless, we observed a shift of myofibers repartition to a smaller CSA sizes in 8-month mice (Fig4D) and this shift was maintained in 12-month-adult mice (Fig4E). The change in myonuclei positioning was not associated with an alteration in the total number of myofibers per muscle (Supplementary Fig3G) or with a massive regeneration process as evidenced by the absence of staining for embryonic myosin heavy chain in myofibers with mispositioned myonuclei (data not shown). We confirmed the accumulation of mislocalized myonuclei in myofibers from 12-month-adult mice, using single isolated myofibers from the *Tibialis Anterior* or *Extensor Digitorum Longus* muscles (Fig4F-G, green arrows). Altogether, these results demonstrate that MACF1 is implicated in the maintenance of myonuclei localization in muscle myofibers of adult mice that is not associated to a regeneration process. This role for MACF1 seems to be a long-term process since no significant myonuclei mis-localization is observed in younger mice.

**Figure 4:**
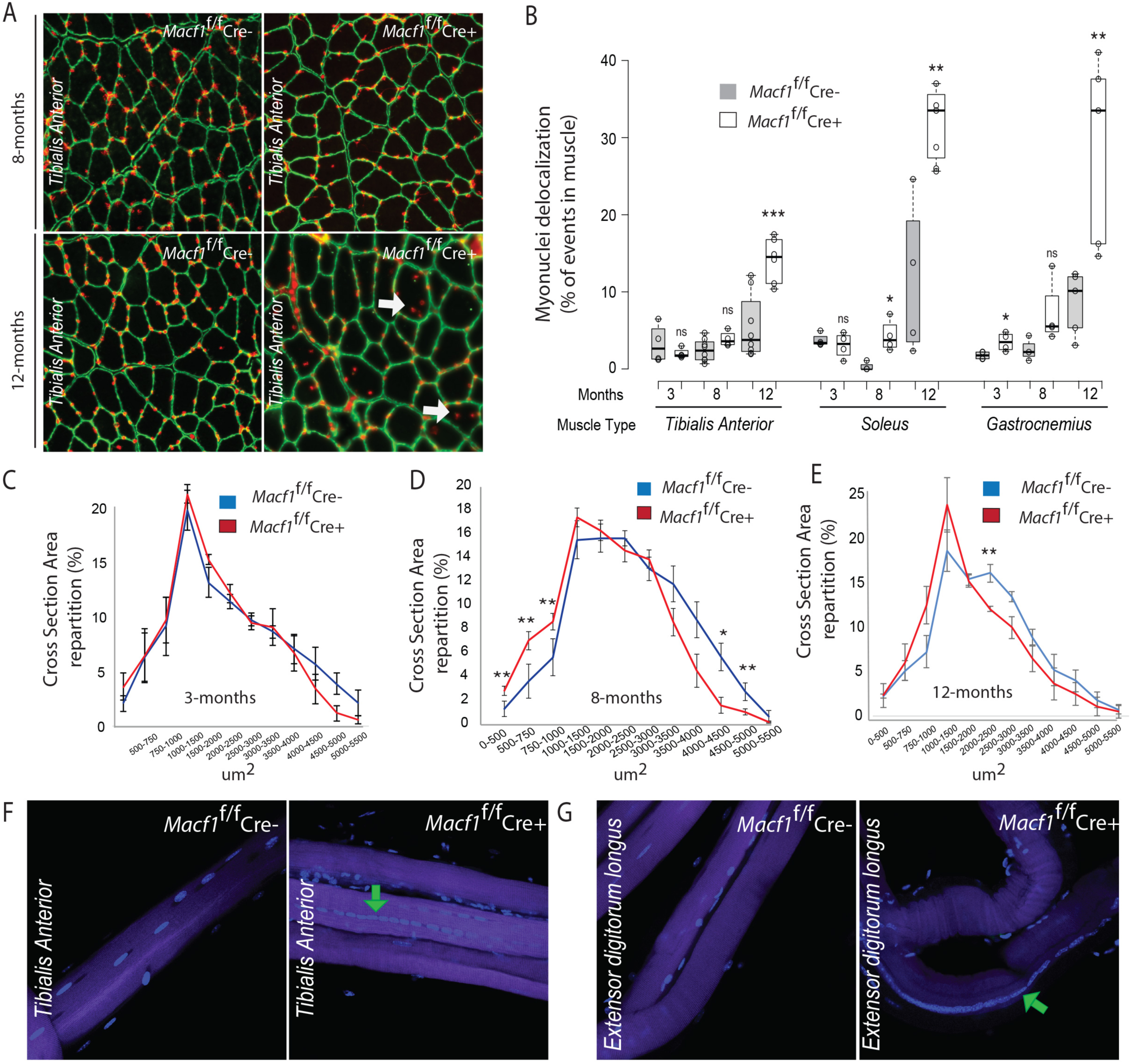
muscle specific MACF1 knockout presents myonuclei organization failure. (A) Representative images of Tibialis Anterior muscle cross-section from *Macf1*^f/f^ Cre- and *Macf1*^f/f^ Cre+ mice at the age of 8 and 12 months-old, stained for Dapi (red) and Laminin (green). Example of myofibers with mis-localized myonuclei are stamped by white arrows. Scale Bar = 150 µm. (B) Quantification of the percentage of myofibers with mis-localized myonuclei in 3, 8 and 12 months-old *Macf1*^f/f^ Cre- and *Macf1*^f/f^ Cre+ in *Tibialis Anterior, Soleus* and *Gastrocnemius* muscles. ***p < 0.001, **p < 0.01, *p < 0.05. Distribution of cross-sectioned myofibers areas from the *Tibialis Anterior* in 3 (C), 8 (D) and 12 (E) months-old *Macf1*^f/f^ Cre- and *Macf1*^f/f^ Cre+ mice. Representative images of *Tibialis Anterior* (F) or *Extensor Digitorum Longus* (G) extracted myofibers stained for Dapi (blue) and Phalloidin (purple) from 12 months-old *Macf1*^f/f^ Cre- and *Macf1*^f/f^ Cre+ mice. Mis-localized myonuclei are stamped by green arrows. At least 3 mice per condition were analyzed. Scale Bar = 150 µm.

### MACF1 muscle-KO mice exhibit early neuromuscular junction alteration with a reset of the related “muscle-microtubule code”

Our *in vitro* results suggested that MACF1 is implicated in myonuclei spreading in an Agrin/NMJ dependent manner (Fig1-2). Thus, we wondered if in our *in vivo* conditional mutant model, synaptic AChRs clustering size and density were impacted by the absence of MACF1. To address this question, we isolated individual’s myofibers from the *Tibialis Anterior* and the *Extensor Digitorum Longus* muscles and visualized AChRs clustering using bungarotoxin staining in 12-months-adult mice (Fig5A). In this condition, we found that the size of synaptic AChRs clusters was highly diminished in both muscles. To determine if the fragmentation of NMJ-related AChRs was an early process, we isolated individual’s myofibers from the *Rectus Lateralis* muscle, known to be multi-innervated, allowing an access to multiple NMJs, from 4-months-young mice (Fig 5B). In this muscle, NMJs were much more fragmented in conditionally mutant mice compared to control mice reflected by the doubling of individual clusters stained for AChRs (Fig5C). Accordingly we measured a reduction of myonuclei associated with the NMJs, with a mean value of 5 myonuclei in control mice (*Macf1*^f/f^ Cre-) as compared to 3 in mutant mice (*Macf1*^f/f^ Cre+) (Fig5D). These data further demonstrate that MACF1 is required to maintain neuromuscular synapses integrity through AChRs clustering in young and adult mice and this alteration seems to precede myonuclei disorganization in myofibers.

**Figure 5:**
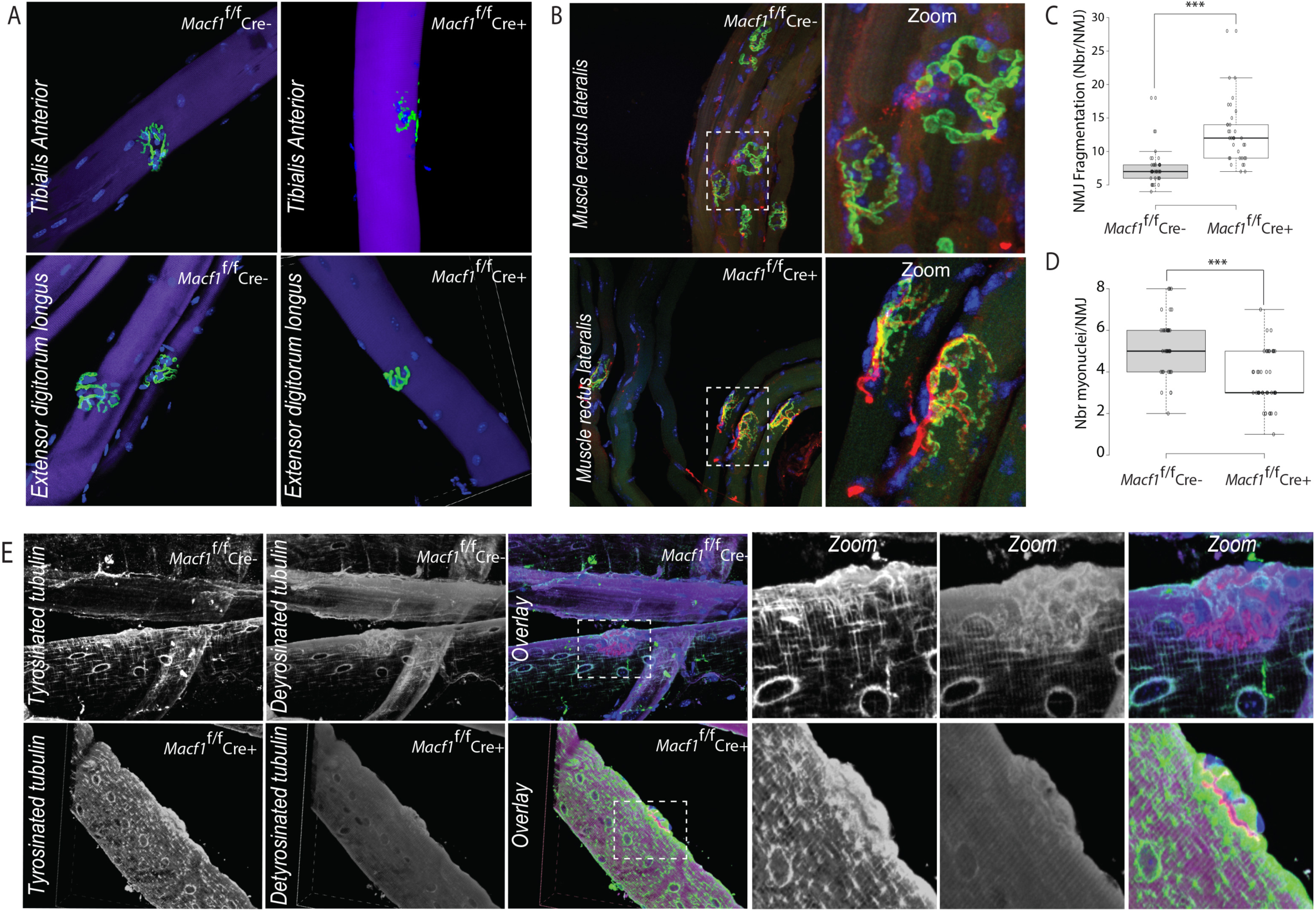
muscle specific MACF1 knockout mice display NMJ integrity failure in parallel with the loss of dynamic microtubules. (A) Representative images of *Tibialis Anterior* (Top) or *Extensor Digitorum Longus* (Bottom) extracted myofibers from 12 months-old *Macf1*^f/f^ Cre- and *Macf1*^f/f^ Cre+ mice stained for Dapi (blue), Phalloidin (purple) and Bungarotoxin (green). Scale Bar = 150 µm. (B) Representative images of extracted myofibers from *Rectus Lateralis* muscle stained for Dapi (blue), presynaptic markers (Neurofilament & Synaptic vesicles 2 (red)) and Bungarotoxin (green) from 4 months-old *Macf1*^f/f^ Cre- and *Macf1*^f/f^ Cre+ mice. Scale Bar = 150 µm. (C) Quantification of NMJ (bungarotoxin staining) fragmentation. (D) Quantification of myonuclei number under Bungarotoxin staining. ***p < 0.001. At least 3 mice per condition were used, n=37. (E) Representative images of *Tibialis Anterior* extracted myofibers stained for Tyrosinated-Tubulin or de-Tyrosinated-Tubulin. In the overlay Tyrosinated-Tubulin (blue), de-Tyrosinated-Tubulin (green) and Bungarotoxin (red) from 12 months-old *Macf1*^f/f^ Cre- and *Macf1*^f/f^ Cre+ mice.

MACF1 contains a microtubule-binding domain (MTB) in the C-terminus part (Supplementary Fig1B) and has been shown to modulate microtubule dynamics through the interaction with different partners such as CLAPS2, EB1, CAMSAP3, Nesprin, Map1B, ErbB2 (Ryan et al., 2012; Ka et al., 2014; Drabek et al., 2006; Noordstra et al., 2016; Wang et al., 2015; Zaoui et al., 2010). In addition, microtubules are subjected to a variety of post-translational modifications (PTMs). A combination of different α- and β-tubulin isoforms and PTMs are referred as the “tubulin code” (Nieuwenhuis and Brummelkamp, 2019). Among these different PTMs, Tubulin de-tyrosination is associated with longer-lived microtubules, whereas more dynamic microtubules are found to be mainly tyrosinated (Bulinski and Gundersen, 1991). To see if MACF1 plays a role in the correct formation of tubulin code, we addressed changes in microtubule tyrosination/de-tyrosination status in conditionally mutant mice compared to control mice (Fig5E). In *Macf1*^f/f^ Cre-muscle, tyrosinated-tubulin was found to locate preferentially at the vicinity of myonuclei along myofibers. No particular enrichment was found close to the AChRs clusters at the NMJ (Fig5E, top). Nonetheless, de-tyrosinated-tubulin was found at the vicinity of myonuclei with a noticeable enrichment at the NMJ, close to the AChRs clusters (Fig5E, top). In conditionally mutant mice (*Macf1*^f/f^ Cre+), although myonuclei shape was changed, tyrosinated-tubulin was still found at the vicinity of myonuclei associated with few aggregates along myofibers, but this time, the presence of tyrosinated-tubulin was enriched at the vicinity of AChRs clusters. In the same perspective, de-tyrosinated-tubulin was highly reduced both around myonuclei and near AChRs clusters (Fig5E, bottom). These data show that MACF1 plays a role in the maintenance of the pool of de-tyrosinated-tubulin which consequently stabilizes the microtubule network at both the vicinity of myonuclei and at the NMJs.

### Adult MACF1 muscle-KO mice exhibit increased myofibers with high mitochondria content

Our *in vitro* results suggested that MACF1 is implicated in mitochondria spreading and fragmentation in muscle fibers (Fig2A, D-E). Since skeletal muscles are composed of a functional and metabolic continuum of slow (type I) and fast fibers (types IIa and IIx), we first questioned whether there were changes in the mitochondrial pool in adult conditional mice. We measured the intensity of succinate dehydrogenase staining, indicative of mitochondrial activity and found a significant increase in the number of fibers with positive succinate dehydrogenase staining in *Tibialis Anterior* muscle in conditionally mutant mice (*Macf1*^f/f^ Cre+) compared to control mice, while no effect was observed in *Soleus* muscle (Fig6A-B). Quantification of proteins comprising the electron transport chain was then performed on *Gastrocnemius* muscles and our results confirmed an increase in total amount of mitochondria (Tom20 relative to actin) and no changes in protein contents from the electron transport chain (CI, II, III, IV and V relative to Tom20) (Fig6C). The intensity of succinate dehydrogenase staining is commonly used to discriminate slow and fast fibers. Since our quantification suggests a change in the repartition of slow/fast fibers in *Macf1*^f/f^ Cre+ muscle, we next investigated if the proportion of slow fibers was changed in conditional mutant mice (*Macf1*^f/f^ Cre+). Cross-sections of 12-month-adult muscles from the *Tibialis Anterior, Soleus* and *Gastrocnemius* were analyzed for slow myosin content and no alteration of the proportion in each muscle type was observed compared to control mice (Fig6D-E). We then used electron microscopy to visualize the ultrastructure of mitochondria in *Tibialis Anterior* muscle of *Macf1*^f/f^ Cre+ mice (Fig6F). This approach confirmed an apparent increase in mitochondria content in muscle fibers associated with the presence of spherical mitochondria in-between myofibrils and close to myonuclei (Fig6F). All together, these data show that MACF1 is involved in the maintenance of mitochondria architecture in muscle fibers and that the loss of MACF1 is associated with a redistribution of mitochondrial content dependent on muscle type.

**Figure 6:**
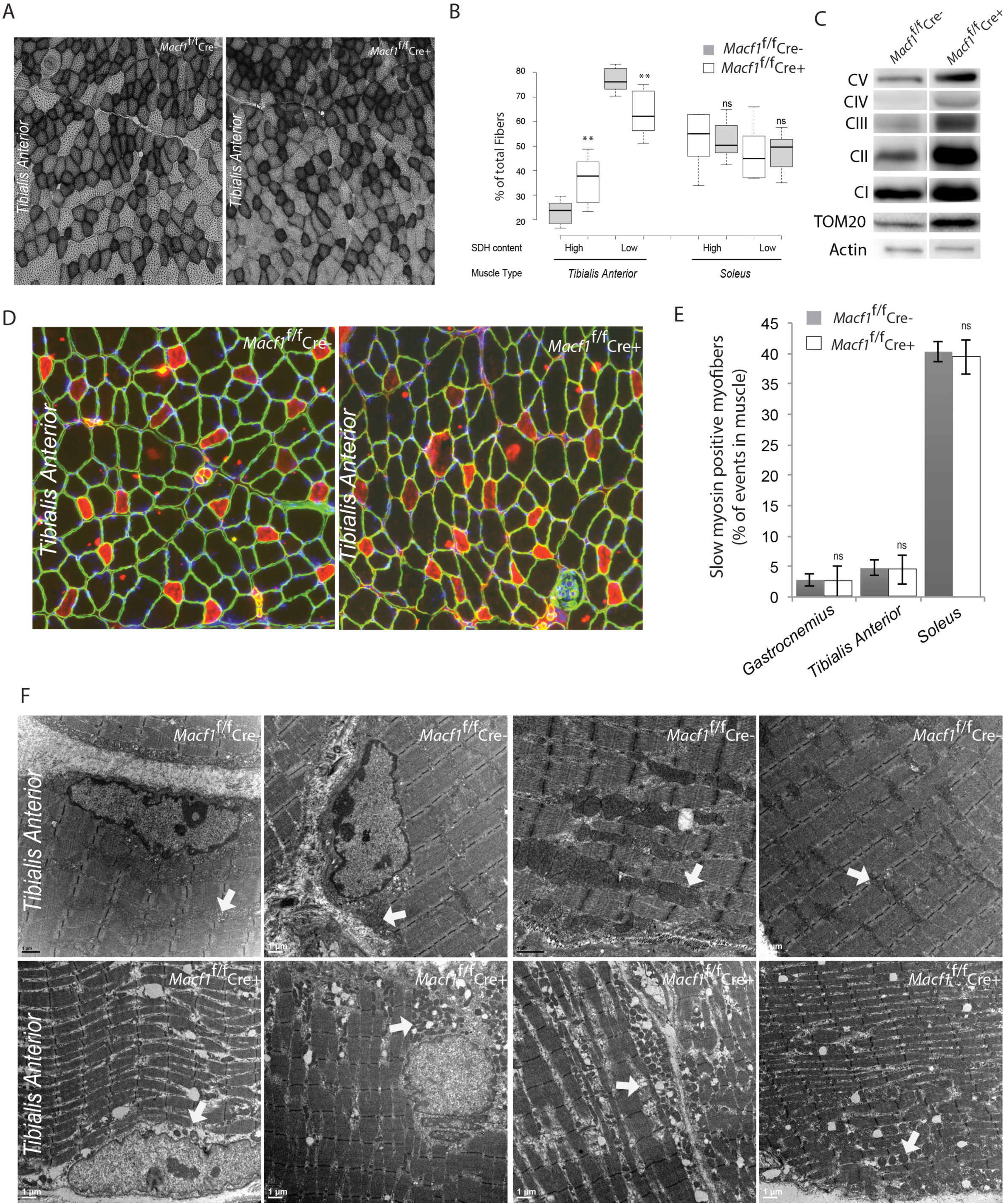
muscle specific MACF1 knockout affects mitochondrial content and structure. (A) Representative images of transversal cross section of *Tibialis Anterior* muscle from 12 months-old *Macf1*^f/f^ Cre- and *Macf1*^f/f^ Cre+ mice stained Succinate DeHydrogenase activity. Scale Bar = 150 µm. (B) Quantification of the Succinate DeHydrogenase activity relative to myofibers repartition in conditional mutant mice compare to control, **p < 0.01. (C) Western blot analysis of OXPHOS proteins, TOM20 and Actin expressions in total *Gastrocnemius* muscles protein extracts from 12 months-old *Macf1*^f/f^ Cre- and *Macf1*^f/f^ Cre+ mice. (D) Representative images of transversal cross section of *Tibialis Anterior* muscle from 12 months-old *Macf1*^f/f^ Cre- and *Macf1*^f/f^ Cre+ mice stained for slow myosin (red) and laminin (green). Scale Bar = 150 µm. (E) Quantification of the percentage of myofibers with slow myosin expression in 12 months-old *Macf1*^f/f^ Cre- and *Macf1*^/f^ Cre+ in *Tibialis Anterior, Soleus* and *Gastrocnemius* muscles. (F) Representative electronic microscopy images of myofibrils and mitochondria organization of *Tibialis Anterior* muscles from 12 months-old *Macf1*^f/f^ Cre- and *Macf1*^f/f^ Cre+ mice. Scale Bar = 1 µm.

### Muscle fibers from MACF1-KO mice show muscle function defects and increased fatigue resistance but preserved excitation-contraction coupling

To see if the observed alterations have any impact on muscle force of our MACF1muscle-KO mice, we examined the muscle performance in *Macf1* conditional KO mutant mice. We used a strictly non-invasive experimental setup offering the possibility to stimulate the Hindlimb muscles to record the force production concomitantly. For both *Macf1*^f/f^ Cre- and *Macf1*^f/f^ Cre+ mice, we assessed the force production from the Hindlimb muscles in response to various incremental stimulation frequencies (i.e., from 1 to 100 Hz) in order to obtain the force-frequency relation. In young 4-months-old mice, we observed a large rightward shift of the force–frequency relation in conditional *Macf1* KO mice (Fig7A), suggesting an alteration of the excitation-contraction coupling process. Interestingly, this shift was maintained in adult mice but in a less significant manner indicating compensation mechanisms to thwart the loss of efficiency in response to motoneurons stimulation. As the absolute maximal force developed by *Macf1* conditional KO mice was not changed compared to control mice (data not shown), we next questioned the fatigability of those muscles. In accordance with the increased SDH staining, we found that in both young and old mice, *Macf1* conditional KO mice are more resistant to fatigue compared to control mice (Fig7B). We then used electron microscopy to study the ultrastructure of muscle in *Macf1*^f/f^ Cre+ mice and analyzed sarcomeres, transverse tubules (T-tubules) and sarcoplasmic reticulum (SR) organization (Fig7C). We found that global muscle organization reflected by sarcomeric structure (myofibrils) is not dependent on MACF1. However, T-tubules and the SR appeared severely affected in some myofibers (Fig7C) confirming a possible direct modification of excitation-contraction coupling in muscle of the *Macf1*^f/f^ Cre+ mice. To test this possibility, we compared voltage-activated SR Ca^2+^ release in *Flexor Digitalis Brevis* (FDB) muscle fibers isolated from conditional *Macf1*^f/f^ Cre+ mice and control mice. Confocal staining using di-8-anepps showed an overall comparable structure of T-tubule network in *Macf1*^f/f^ Cre- and in *Macf1*^f/f^ Cre+ FDB muscle fibers (Fig7D-top, & -middle panels). However, some isolated muscle fibers exhibited strong alteration of T-tubule orientation without perturbation in T-tubules density or sarcomere length (Fig7D-Bottom panel, E-G). We next tested if SR Ca^2+^ release amplitude and kinetics are affected in *Macf1*^f/f^ Cre+ muscles fibers. Figure 7H shows rhod-2 Ca^2+^ transients elicited by membrane depolarizing steps of increasing amplitude in a *Macf1*^f/f^ Cre- and in a *Macf1*^f/f^ Cre+ fibers. As routinely observed under these conditions (Kutchukian et al., 2017), transients in control fibers exhibit a fast early rising phase upon depolarization followed by a slower phase at low and intermediate voltages and by slowly decaying phase for the largest depolarizing steps. As shown in figure 7I, rhod-2 transients from *Macf1*^f/f^ Cre+ fibers exhibit an overall similar time-course, not distinguishable from control muscle fibers. In each tested fiber, the rate of SR Ca^2+^ release was calculated from the rhod-2 Ca^2+^ transients. Traces for the calculated rate of SR Ca^2+^ release corresponding to the transients shown in figure 7I are shown in figure 7J. In both fibers, the rate exhibits a similar early peak, the amplitude of which increases with the amplitude of the pulse, followed by a spontaneous decay down to a low level. The SR Ca^2+^ release peak amplitude was similar in *Macf1*^f/f^ Cre- and in a *Macf1*^f/f^ Cre+ muscle fibers for all voltages (Fig7J) and the time to reach the peak was also not affected in the *Macf1*^f/f^ Cre+ muscle fibers (Fig.7K, values not shown). Mean values for maximal rate of SR Ca^2+^ release (Max d[Ca_tot_]/dt), mid-activation voltage (V_0.5_) and slope factor (k) of the voltage-dependence were statistically unchanged in *Macf1*^f/f^ Cre+ muscle fibers compared to control fibers (Fig7K). Accordingly, there was also no change in the voltage-dependent Ca^2+^ channel activity of the dihydropyridine receptor (also referred to as Ca_V_1.1, the voltage-sensor of excitation-contraction coupling) in fibers from *Macf1*^f/f^ Cre+ mice (not shown). All in all, these data show that muscle from *Macf1* conditional mutant mice exhibit delay in excitability in response to nerve stimulation and although the T-tubule network organization seems affected in some myofibers, the global efficiency of SR-Ca^2+^ release seems not engaged. Interestingly, muscles from *Macf1* conditional mice display a higher resistance to fatigue compared to muscles of control mice.

**Figure 7:**
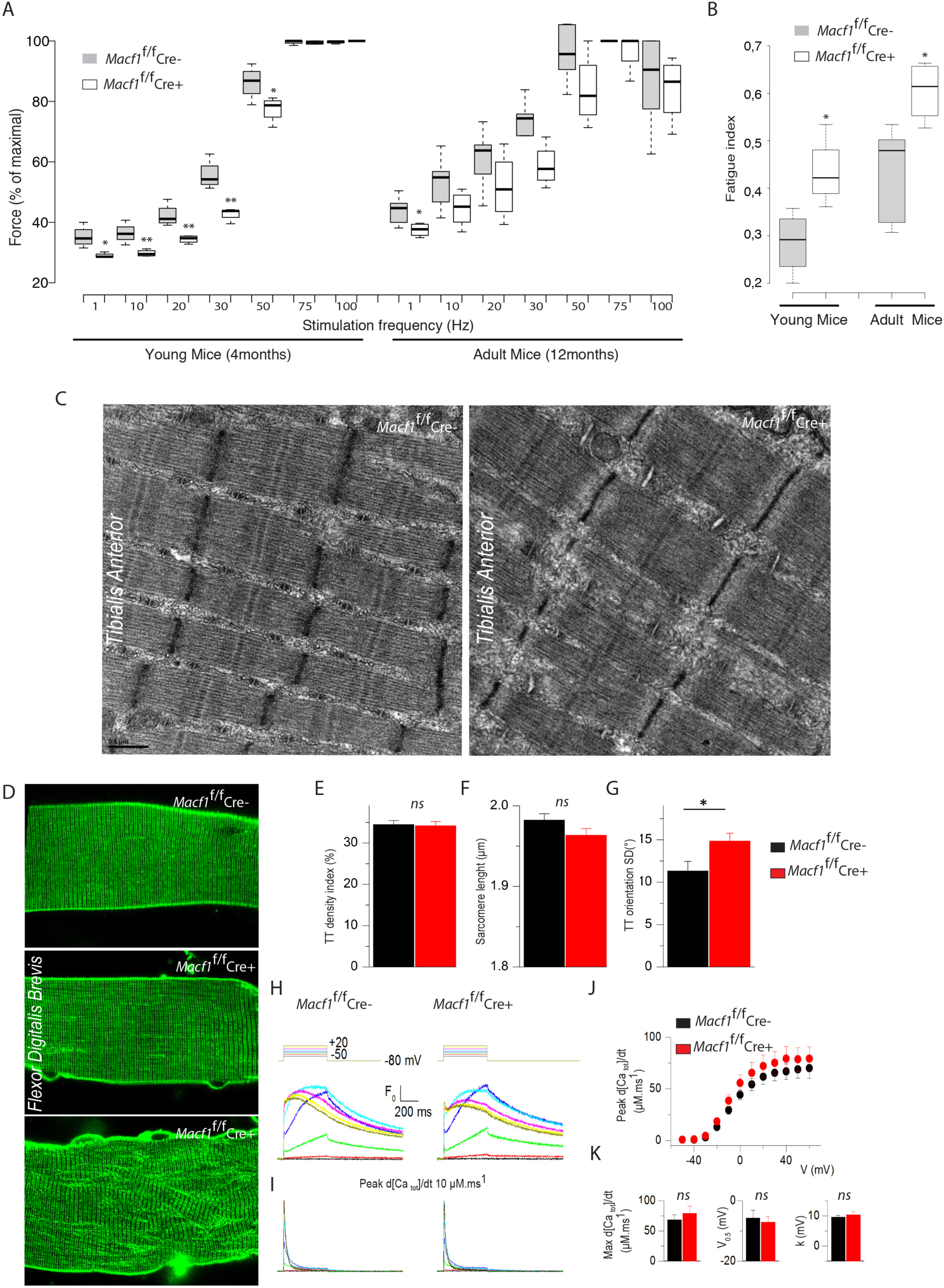
muscle specific *Macf1* knockout affects muscle functions and T-tubule formation but not RS Ca2^+^ release. (A) Force production from the *Hindlimb* muscles in young (4 months) and adult (12 months) *Macf1*^f/f^ Cre- and *Macf1*^f/f^ Cre+ mice in response to various incremental stimulation frequencies (from 1 to 100 Hz). **p < 0.01, *p < 0.05. (B) Quantification of the fatigue index from the *Hindlimb* muscles in young (4 months) and adult (12 months) *Macf1*^f/f^ Cre- and *Macf1*^f/f^ Cre+. *p < 0.05. (C) Representative electronic microscopy images of myofibrils and T-tubule organization of *Tibialis Anterior* muscles from 12 months-old *Macf1*^f/f^ Cre- and *Macf1*^f/f^ Cre+ mice. Scale Bar = 0,5 µm. (D) Representatives fluorescence images of di-8-anepps staining of the T-tubule network in a *Macf1*^f/f^ Cre- and *Macf1*^f/f^ Cre+ mice muscle fibers. Quantification of T-tubule density index (E), Sarcomere length (F) and T-Tubules orientation (G) in a *Macf1*^f/f^ Cre- and *Macf1*^f/f^ Cre+ mice muscle fibers. (H) Representative rhod-2 Ca^2+^ transients in a *Macf1*^f/f^ Cre- and in a *Macf1*^f/f^ Cre+ fiber in response to 0.5s long depolarizing pulses from −80mV to the range of indicated values, with intervals of 10 mV. (I) Corresponding Ca^2+^ release flux (d[CaTot]/dt) traces calculated as described in the Methods. (J) Mean voltage-dependence of the peak rate of SR Ca^2+^ release in *Macf1*^f/f^ Cre- and in a *Macf1*^f/f^ Cre+ fibers. (K) Inset shows the mean values for maximal rate, half velocity and k index of SR Ca^2+^ release in the two groups of fibers, as assessed from Boltzmann fits to data from each fiber.

## Discussion

Different mechanisms control myonuclei positioning during muscle fiber formation (Roman and Gomes, 2017; Starr, 2017; Manhart et al., 2018). Early steps of myonuclei positioning in immature fibers (myotubes) depend on an interplay between microtubule, microtubule-associated-proteins and motors proteins such as MAP4, MAP7, Dynein and Kif5b (Gimpel et al., 2017; Mogessie et al., 2015; Metzger et al., 2012; Cadot et al., 2012; Wang et al., 2013; Wilson and Holzbaur, 2014). This process also depends on myonuclei membrane associated components such as Nesprin-1 or AKAP9 (Wilson and Holzbaur, 2015; Gimpel et al., 2017). In mature myofibers, myonuclei are found at the periphery of fibers and reach this localization by a process dependent on actin network, contraction and myonuclei rigidity (Roman et al., 2017). However, processes involved in maintenance of myonuclei at the periphery of muscle fibers are still unknown.

The present study is the first to demonstrate that a protein related to the microtubule network controls and maintains peripheral myonuclei localization in mature myofibers. Here, we show that MACF1 is progressively accumulated at the onset of myotubes formation, with no evident implication in early steps of myonuclei spreading or myotubes elongation but remarkable actions in maturation steps to set the distance between adjacent myonuclei (Fig1-2). Accordingly, conditionally mutant mice (*Macf1*^f/f^ Cre+) exhibit a strong decrease in peripheral myonuclei, but surprisingly, this alteration becomes significant only at 12-months of age (Fig4). This alteration comes after a slight reduction in myonuclear domains in myofibers reflected by a decrease in the mean diameter of myofibers in 8-month-adult mice. These data suggest a complex interplay between factors that maintain peripheral myonuclei localization and demonstrate that the absence of MACF1 contributes to a long-term modification of this equilibrium that breaks in 12 month-adult *Macf1*^f/f^ Cre+ mice.

It has already been shown that atrophy can result from a progressive alteration of NMJs (Bonaldo and Sandri, 2013). In fact, in our *in vivo* model, the first alterations in myonuclei localization occurred at the NMJs in 4-month-adult mice (Fig5). This is in accordance with results from our *in vitro* system, where myonuclei disruption in myofibers occurs when myofibers are stimulated with extra-cellular matrix and neural Agrin, which seems to be the starting point of myonuclei positioning alteration in myofibers. Myonuclei clustering beneath the NMJ is known to be dependent on proteins of the LINC complex such as Nesprin and Sun proteins and implicated myonuclei rigidity, as mutation or depletion of Lamin A/C leads also to aberrant and fragile NMJs with myonuclei lying out of it (Razafsky and Hodzic, 2015; Méjat et al., 2009). In the present study, we show that MACF1 contributes in a first instance to synaptic myonuclei clustering at the NMJs. This property seems to be related to the control of MACF1 on microtubule dynamics and stability (Fig3 & 5). Microtubule dynamics and associated MAPs (Claps2, Clip170) and motors (Dynein) mediate microtubule capture at the vicinity of synaptic myonuclei and seem to be crucial for the stabilization of acetylcholine receptors (AChRs) and Musk (muscle-specific receptor tyrosine kinase) at the post-synaptic region (Basu et al., 2015; Vilmont et al., 2016; Schmidt et al., 2012; Basu et al., 2014). Here, we found that MACF1 controls myonuclei-related “microtubule code” affecting the ratio between tyrosinated and non-tyrosinated tubulin and thus longer-lived microtubules at the vicinity of myonuclei (Bulinski and Gundersen, 1991). Interestingly, our observations point to a strong influence of MACF1 at axons endplates to control myonuclei clustering and microtubule organization dynamics and this make sense as this site is known to strongly accumulates microtubule network (Ralston et al., 2001). Synaptic alteration in the local “microtubule code” can itself drive changes in forces applied locally on synaptic myonuclei, independently of myonuclei membrane proteins, and contribute to the NMJ integrity. Interestingly, extra-synaptic myonuclei are affected by this change of dynamics after the synaptic myonuclei, suggesting a different pool of proteins controlling myonuclei localization between synaptic and extra-synaptic myonuclei.

Genetic inactivation of *Macf1* redistributes microtubule dynamic equilibrium, which acts on diverse pathways that control muscle functionality. MACF1 directly affects NMJ integrity through the control of AChRs clustering (Fig2 & 4). We observe an early failure in the capacity to cluster AChRs *in vitro* in mature myofibers (Fig2). This effect is confirmed in conditionally mutant mice (*Macf1*^f/f^ Cre+), which exhibit failure in AChRs clustering at the contact with motoneurons. This specific alteration of NMJ integrity is most likely responsible for the decrease in muscle performance (Fig5). This possibility is reinforced by the facts that first, myofibril genesis is not impacted either *in vitro* in mature *Macf1*-KD myofibers or *in vivo* in mutant mice (*Macf1*^f/f^ Cre+) and second, there is no alteration in SR-Ca^2+^ release even though the overall structure of T-tubule network seems to be affected in few myofibers (Fig7). In contrast, we observed, in our *in vitro* model, an alteration in the organization of the mitochondrial network (Fig2). This modification is translated in the conditional mutant mice into a strong increase of mitochondria with shape alteration, mainly smaller and rounder (Fig6). Interestingly, we also observe an increase in the number of oxidative fibers by muscle and it appears to be dependent on muscle type, without changes in myofibers typology according to myosin type expression (fast or slow muscle fibers). This alteration in oxidative fibers content is correlated with an increase of the resistance to fatigue (Fig6). This effect of MACF1 down-expression on mitochondria organization is consistent with a number of previous observations, namely, microtubule network, through molecular motors such as Kif5b and Dynein, previously shown to control mitochondrial movements in myoblasts (iqbal and Hood, 2014). Additionally, MACF1 was shown to affect mitochondria localization through nuclear positioning in oocyte egg (Escobar-Aguirre et al., 2017). Interestingly, the interplay between actin branching and microtubule acetylation has been recently pointed out as a mitochondrial distribution keeper in fibroblasts cells (Shi et al., 2019). Increase in microtubule dynamic run against stabilization of microtubules partners, but even worse, it can contribute to the delocalization of few MAPs that preferentially bind to stable microtubule (Gache et al., 2010). Among them, DNM2, is a MAP that preferentially associate with stable microtubules (Tanabe and Takei, 2009). The absence of MACF1 could contribute to the release of DNM2 clustered on stable microtubule and consequently trigger mitochondria fission in mutant mice (*Macf1*^f/f^ Cre+) (Kraus and Ryan, 2017). Alternatively, increase in microtubule dynamic instability directly impact local concentration of free tubulin (Gache et al., 2005) whereas free available tubulin-βII was shown to affect mitochondrial metabolism and organization (Kumazawa et al., 2014; Guzun et al., 2011; Guerrero et al., 2009). In this perspective, MACF1 could command mitochondria shape and functionality through the control of microtubule dynamics in muscle fibers.

Various diseases can directly affect NMJs integrity, resulting in a progressive disconnection between muscles and motoneurons. Two of the most common laminopathies, Limb-Girdle Muscular Dystrophy (LGMD) and Emery-Dreifuss Muscular Dystrophy (EDMD), which affect striated skeletal muscles, typically lead to a disproportion of fiber size, myonuclei aggregation along myofibers and dispersion under NMJs (Chojnowski et al., 2015). Conversely pathological mechanisms affecting presynaptic site can similarly lead to synaptic myonuclei dispersion as is the case in MotoNeuron Diseases (MNDs), such as Amyotrophic Lateral Sclerosis (ALS), a fatal neurodegenerative condition characterized by progressive motoneuron denervation and muscle atrophy associated with synaptic myonuclei dispersion (Geevasinga et al., 2016). Additionally, there are several conditions where peripheral myonuclei localization is affected. For example, in a heterogeneous group of inherited muscle diseases, CentroNuclear Myopathies (CNMs), are defined pathologically by an abnormal localization of myonuclei in the center of muscle fibers (Jungbluth and Gautel, 2014). But unlike many myopathies, CNMs are not linked to excessive degeneration/regeneration processes. Genes implicated in various forms of CNMs as well as encode proteins that participate in different aspect of membrane remodeling and trafficking such as Mtm1, Amphyphisin-2/BIN1, Dynamin-2 and Hacd1have been already identified (Laporte et al., 1996; Bitoun et al., 2005; Muller et al., 2003; Muhammad et al., 2013). In addition, in the case of Sarcopenia, peripheral myonuclei are also progressively lost (Edström et al., 2007; Malatesta et al., 2009; Brooks et al., 2009). Altogether, MACF1 will be a perfect candidate, from the postsynaptic muscle view, to investigate the basis of motoneuron diseases and the “de-peripherization” of myonuclei in skeletal muscle in various types of myopathies.

---

**Supplementary figure 1:**
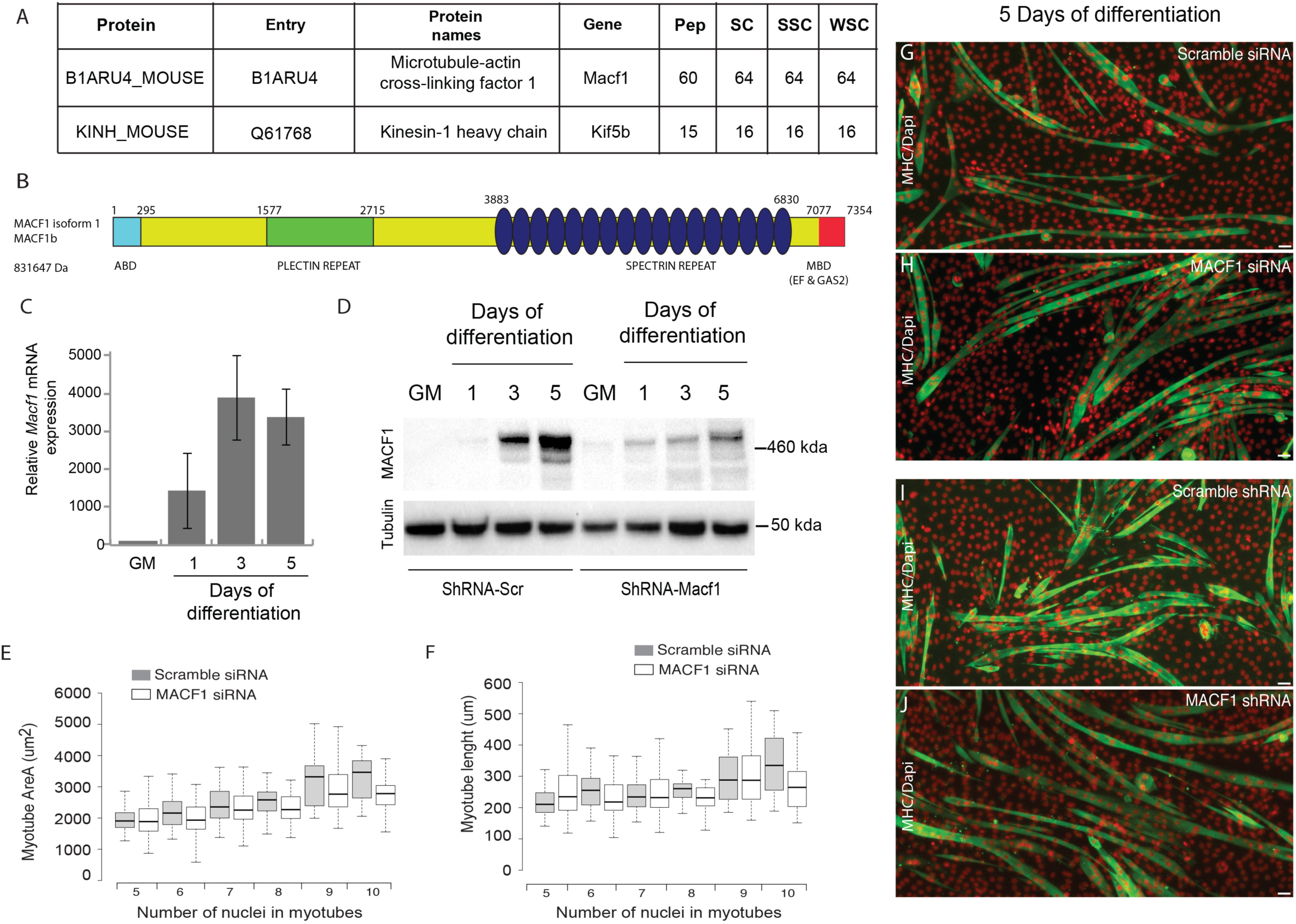
MACF1 expression in myotubes and myofibers. (A) Table showing mass spectrometry identification of MACF1 and Kif4b in 3 days-old primary myotubes. (B) Schematic representation of MACF1 already identified domains. ABD is the “Actin Binding Domain” and MBD is the “Microtubule Binding Domain”. (C) QRT-PCR analysis of *Macf1* gene expression level relative to *Hprt1* and *Gapdh* gene in proliferating C2C12 cells (GM) and after 1, 3, or 5 days of differentiation. Data’s means are ± SEM (n=3). (D) Western blot analysis of MACF1 protein expression in total protein extracts from proliferating cells (GM) to differentiating C2C12 myotubes after 1, 3, or 5 days of differentiation in cells treated with either Scramble shRNA (control) or a pool of 4 individual shRNAs targeting *Macf1* gene. Tubulin is used as “loading control”. Myotubes Area (E) and myotubes length (F) ranked by myonuclei content per myotubes were quantified after 3 days of differentiation in cells treated with a scramble or a pool of 3 individual *Macf1* siRNAs. In E-H, 3 individual experiments, n=170 for scramble and n=365 for SiRNA *Macf1* conditions. (G-J) Immunofluorescence staining of Myosin Heavy Chain (green) and myonuclei (red) in 5-days old C2C12 myotubes treated with scramble siRNA (G) or a pool of 3 individual siRNAs targeting *Macf1* (H) and in C2C12 myotubes treated with scramble shRNA (I) or a pool of 4 individual shRNAs targeting *Macf1* (J). Scale Bar = 50 µm.

**Supplementary figure 2:**
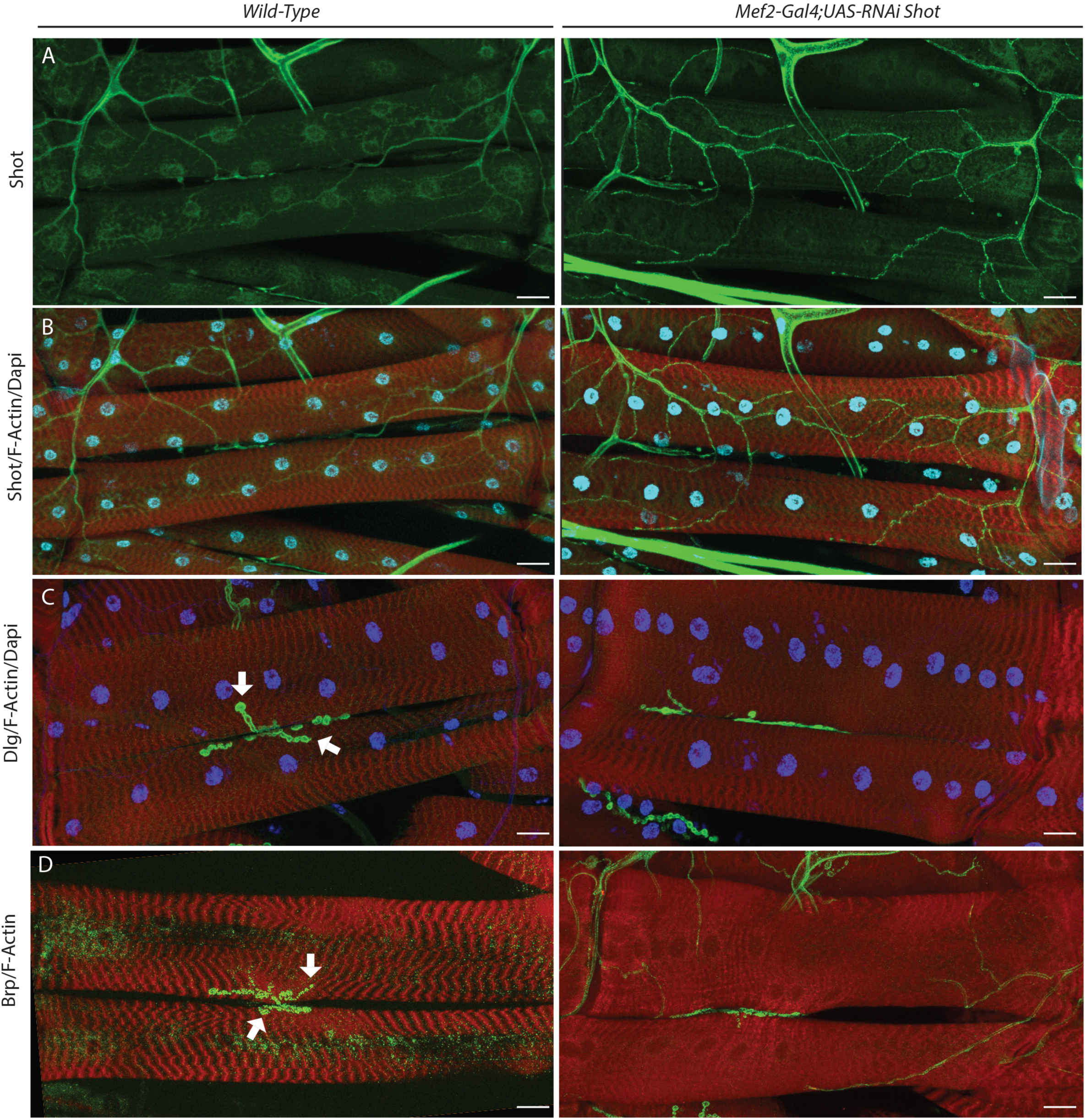
MACF1/Shot in myotubes and myofibers. (A-B) Larval muscles labeled with Shot (Green), Dapi (blue) and Phalloidin (red) in WT (left) or RNAi treated muscle-targeting *Shot* (right panel). (C) Larval muscles labeled with Discs-Large, Dlg (Green), Dapi (blue) and Phalloidin (red) in WT (left) or RNAi treated muscle targeting *Shot* (right panel). (D) Larval muscles labeled with Bruchpilot, BRP (Green), Dapi (blue) and Phalloidin (red) in WT (left) or RNAi treated muscle targeting *Shot* (right panel). Scale Bar = 50 µm.

**Supplementary figure 3:**
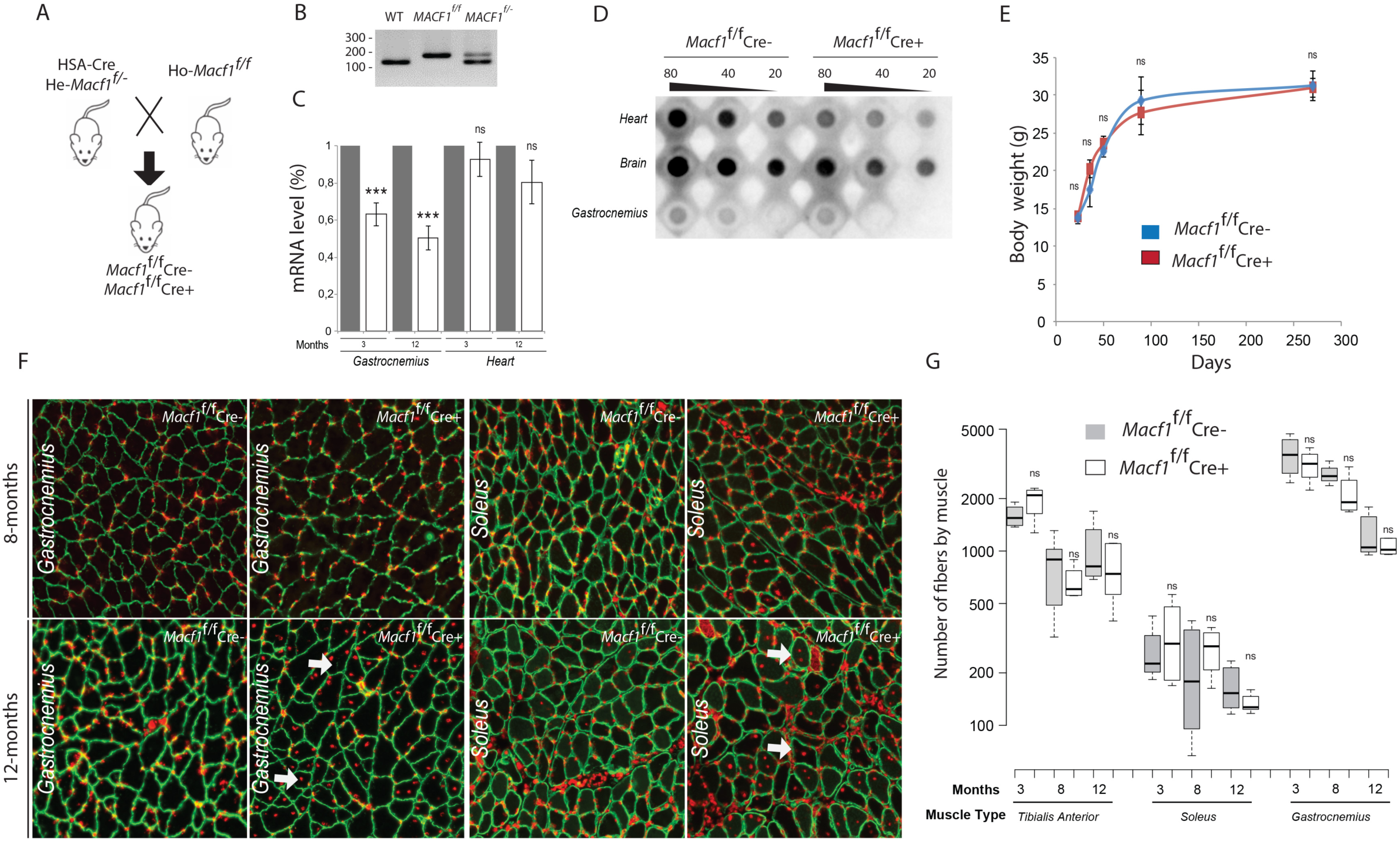
Muscle specific *Macf1* knockout mice. (A) Schema of the strategy used to generate mouse muscle specific *Macf1* knockout mouse model (B) Mutant mice carrying a Cre transgene under the control of the Human Skeletal Actin (HSA) promoter confirmed by genomic PCR. (C) Total MACF1 expression is reduced by Half-fold in *Gastrocnemius* muscle lysates. Mean ± SEM values from three independent experiments. ***p < 0.001. (D) Dot blot assay for MACF1 expression in Heart, Brain and *Gastrocnemius* lysates from *Macf1*^/f^ Cre- and *Macf1*^f/f^ Cre+ (E) Evolution of *Macf1*^f/f^ Cre- and *Macf1*^f/f^ Cre+ body mass of mice during 12 month (n=4). (F) Representative images of *Gastrocnemius* and *Soleus* muscle cross-section in 8 and 12 months-old *Macf1*^f/f^ Cre- and *Macf1*^f/^ Cre+ mice ^f^ stained for Dapi (red) and Laminin (green). Example of myofibers with mis-localized myonuclei are stamped by white arrow. Scale Bar = 150 µm. (G) Quantification of the total number of myofibers in 3, 8 and 12 months-old *Macf1*^f/f^ Cre- and *Macf1*^f/f^ Cre+ mice from in *Tibialis Anterior, Soleus* and *Gastrocnemius muscles.*

## Video legend

**Movie 1.** EB1 comets velocity in primary myotubes transfected with EB1-GFP and scrambled siRNA. Picture were recorded every 500 ms. Movie correspond to Figure 3A Scramble.

**Movie 2.** EB1 comets velocity in primary myotubes transfected with EB1-GFP and a pool of 3 individual siRNAs (#952, #954, #958) targeting *Macf1*. Picture were recorded every 500 ms. Movie correspond to Figure 3A MACF1 siRNA.

**Movie 3.** Nuclear movements in primary myotubes transfected with a scrambled shRNA (green) and lamin-chromobody® (red). Picture were recorded every 15 min. Movie correspond to Figure 3C sh-Scramble.

**Movie 4.** Nuclear movements in primary myotubes transfected with a pool of 4 individual shRNAs targeting *Macf1* (green) and lamin-chromobody® (red). Picture were recorded every 15 min. Movie correspond to Figure 3C sh-MACF1.

## Materials and methods

### Cell culture

Primary myoblasts were collected from wild type C57BL6 mice as described before (Falcone et al., 2014; Pimentel et al., 2017). Briefly, Hindlimb muscles from 6 days pups were extracted and digested with collagenase (Sigma, C9263-1G) and dispase (Roche, 04942078001). After a pre-plating step to discard contaminant cells such as fibroblasts, myoblasts were cultured on matrigel coated-dish (Corning, 356231) and induced to differentiate in myotubes for 2-3 days in differentiation media (DM: IMDM (Gibco, 21980-032) + 2% of horse serum (Gibco, 16050-122) + 1% penicillin-streptomycin (Gibco, 15140-122)). Myotubes were then covered by a concentrated layer of matrigel and maintained for up to 10 days in long differentiation culture medium (LDM: IMDM (Gibco, 21980-032) + 2% of horse serum (Gibco, 16050-122) + 0.1% Agrin + 1% penicillin-streptomycin (Gibco, 15140-122)) until the formation of mature and contracting myofibers. LDM was changed every two days.

Mouse myoblast C2C12 cells were cultured in Dulbecco’s modified Eagle’s medium (DMEM (Gibco, 41966029) + 15% fetal bovine serum (FBS) (Gibco, 10270-106) + 1% penicillin-streptomycin (Gibco, 15140-122))) and were plated on 0.1% matrigel-coated dishes for 1-2 days before differentiation. Differentiation was induced by switching to differentiation media (DMEM + 1% horse serum).

### Production of wild type Agrin recombinant proteins

For production of recombinant proteins, the stably transfected HEK293-EBNA cell lines were grown to about 80 % confluence and were transferred to expression medium without FBS. Conditioned medium containing secreted proteins was collected every 3 days and replaced with fresh expression media for 12 days. Conditioned medium was centrifuged at 2,000xg for 10 min to pellet the cells before storing at -20°C. After thawing, recombinant proteins were purified from conditioned media by HPLC, on a HiTRAP Imac HP column (GE Healthcare, 17-0920-03,), eluted with imidazol and desalted on silica column with PBS (HiPrep 26 /10Desalting, GE Healthcare) or on a Vivaspin column (Vivaspin Sartorius). The absolute concentration of soluble Agrins was estimated on a coomassie blue stained gel by comparison with a known amount of a commercial purified Agrin.

### Cell transfection

For C2C12 cells, 3 different siRNAs Silencer per gene were transfected in cells using Lipofectamine 2000 (ThermoFisher Scientifics, 11668-019) at the final concentration of 10 nM, following manufacturer instructions, 2 days before differentiation. For shRNA cDNA (Geneocopia), we used Lipofectamine 2000 (ThermoFisher Scientifics, 11668-019) was used following manufacturer instructions as well.

For primaries cells, siRNA were transfected using Lipofectamine 2000 (ThermoFisher Scientifics, 11668-019) at the final concentration of 2nM. shRNA (Geneocopia), Eb1 or RFP-Lamin-chromobody (Chromotek) cDNA were transfected in cells using Llipofectamine 3000 (ThermoFisher Scientifics, L3000-008).

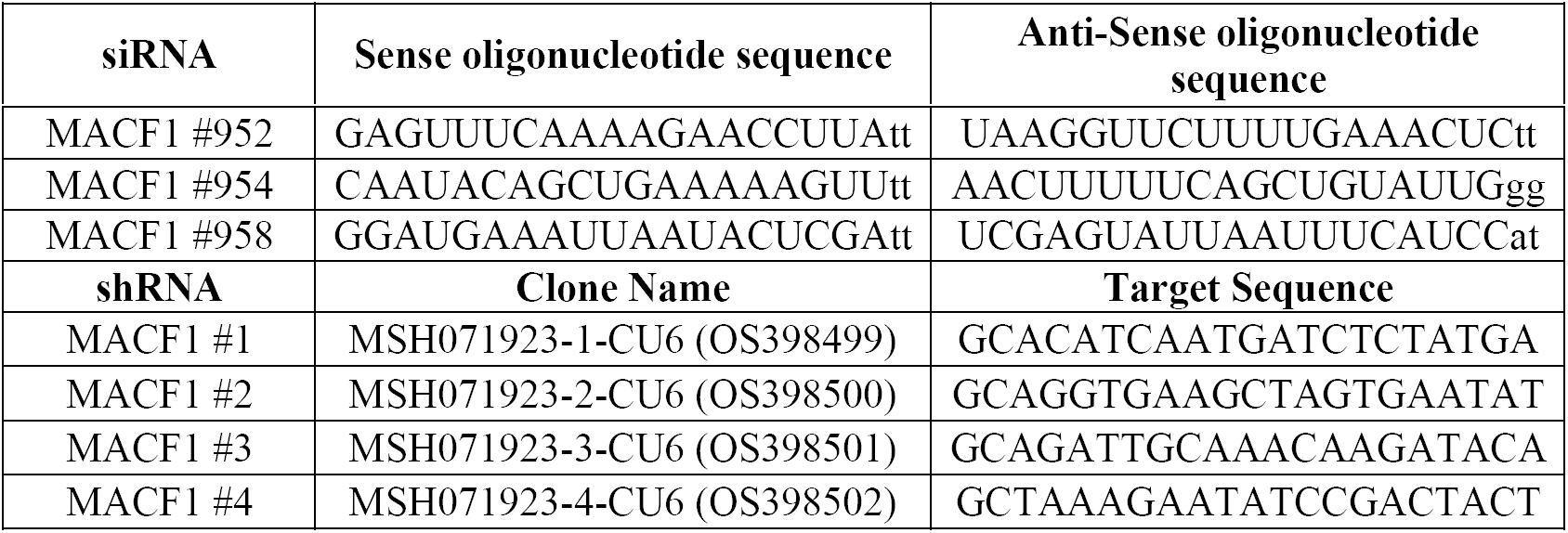

### Protein sample preparation

For primary cultured cells or C2C12 cell lines, cells were harvested, using 1X Trypsin for 5min at 37°C and centrifuged at 1500RPM for 5min at 4°C. Cell pellets were diluted and incubated in the optimal volume of RIPA lysis buffer containing phosphatases inhibitors (Sigma, P5726-5mL) and proteases inhibitors (Sigma, P8340) for 10min at 4°C. Following a sonication and a centrifugation at 12000RPM for 10min at 4°C, protein samples were collected for further uses. The concentration of proteins was determined using BCA protein assay kit (Thermo Fisher Scientifics, 23225) as described by the manufacturer.

### Western blot and dot blot analysis

To carry out western blots, the same amount of sample were loaded in 6% acrylamide gels and were migrated at 130V for 10min followed by 160V for 90min. iBlot 2 mini slacks (Thermo Fisher Scientifics, IB23002) semi-dry system was used to transfer the proteins to nitrocellulose membranes. Membranes were then saturated in 5% milk in TBS for 1H at RT and were incubated in primary antibodies in 5% milk in TBS over night at 4°C. Following washes by 0.1% Tween-20-1X TBS, the membranes were incubated in HRP conjugated secondary antibodies in 5% milk in TBST for 1H at RT. Following washes by 0.1% Tween-20-1X TBS the detection of the target proteins was carried out using Super Signal West Femto (Thermo Fisher Scientifics, 34095) and ChemiDoc imaging system (BioRad).

To carry out dot blots, Bio-Dot SF Microfiltration Apparatus plates (BioRad) were used to transfer the protein samples onto the nitrocellulose membranes. Membranes were then saturated in 5% milk in TBS for 1H at RT and were incubated in primary antibodies over night at 4°C. Following washes by 0.1% Tween-20-1X TBS, the membranes were incubated in HRP conjugated secondary antibodies in 5% milk in TBST for 1H at RT. Following washes by 0.1% Tween-20 in TBS the detection of the target proteins was carried out using Super Signal West Femto (Thermo Fisher Scientifics, 34095) and ChemiDoc imaging system (BioRad).

### Primary cells immunofluorescence staining

Cells were fixed in 4%PFA in PBS for 20min at 37°C followed by washes with PBS and permeabilization with 0.5% Triton-X100 in PBS for 5min at RT. Following washes with PBS, cells were saturated with 1% BSA in PBS for 30min at 37°C and incubated in primary antibodies over night at 4°C. Following washes with 0.05% Triton-X100-1X PBS, cells were incubated in secondary antibodies or dyes for 2H at RT followed by washes with 0.05% Triton-X100 in PBS and a last wash in PBS. Cultured myofibers were imaged using either Z1-AxioObserver (Zeiss) or confocal SP5 microscope (Leica).

### Isolation of mono-myofibers and immunofluorescence staining

Following the dissection of the whole muscle from the mice, TA or EDL muscle blocks were fixed in 4%PFA in PBS for 2H at RT. After washes, 30 to 50 mono-myofibers were isolated per staining from each muscle. Myofibers were then permeabilized using 0.5% Triton-X100 in PBS for 5min at RT and saturated in 1% BSA in PBS for 30min at RT. Based on the experiments, myofibers were incubated in desired primary antibodies at 4°C over night. Following washes with 0.05% Triton-X100 in PBS, myofibers were incubated in secondary antibodies or dyes for 2H at RT followed by washes with 0.05% Triton-X100 in PBS and a last wash in PBS. Myofibers were mounted on slides using fluromount Aqueous mounting (Sigma, F4680-25mL) and kept at 4°C or -20°C. Slides were analyzed using confocal SP5 microscope (Leica) or TI-Eclipse (Nikon).

### Antibodies

| Antibody or Dye | Species and Utility | Dilution Factor | Manufacturer and Reference |
| --- | --- | --- | --- |
| Anti-Cytochrome-C | Mouse - primary | 1:300 | Cell Signaling Technologies 12963S |
| Anti-MACF1 | Rabbit - primary | 1:100 | Santa Cruz SC-68430 |
| Anti-TOM20 | Rabbit - primary | 1:1000 | Cell Signaling Technologies D8T4N |
| Anti-OXPHOS | Mouse - primary | 1:1000 | Abcam Ab110413 |
| Anti-α-Tubulin | Mouse - primary | 1:1000 | Sigma T6074-200uL |
| Anti-HSP90 | Mouse - primary | 1:1000 | Enzo life SPA830 |
| Anti-Tyrosinated Tubulin | Rat - primary | 1:100 | Andrieux's Lab gift |
| Anti-deTyrosinated Tubulin | Rabbit - primary | 1:100 | Andrieux's Lab gift |
| MF20 | Mouse-primary | 1:10 | DSHB |
| Dapi-brilliant blue | Dye | 1:50000 | Thermo Fisher Scientifics D1306 |
| α-Bungarotoxin-Alexa Flour 488 | Dye | 1:1000 | Thermo Fisher Scientifics B13422 |
| α-Bungarotoxin-Alexa Flour 555 | Dye | 1:200 | Thermo Fisher Scientifics B35451 |
| Phalloidin-Alexa Flour 647 | Dye | 1:100 | Thermo Fisher Scientifics A22287 |
| Anti-mouse-HRP | Goat - secondary | 1:3000 | Invitrogen 62-6520 |
| Anti-rabbit-HRP | Goat - secondary | 1:3000 | Invitrogen 65-6120 |
| Anti-rat-Alexa Flour 488 | Donkey - secondary | 1:500 | Thermo Fisher Scientifics A-21208 |
| Anti-rabbit-Alexa Flour 647 | Goat - secondary | 1:500 | Thermo Fisher Scientifics A-21245 |
| Anti-mouse-Alexa Flour 647 | Goat - secondary | 1:500 | Thermo Fisher Scientifics<br>A-21240 |

### Video-Microscopy

Time-lapse images were acquired using Z1-AxioObserver (Zeiss) with intervals of 15minutes. Final videos were analyzed using Metamorph (Zeiss) and SkyPad plugin as described before (Cadot et al., 2014).

### Electronic-Microscopy

Tissues were cut into small pieces and fixed in 2% glutaraldehyde for 2H at 4°C. Samples were washed three times for 1H at 4°C and post-fixed with 2% OsO4 1H at 4°C. Then tissues were dehydrated with an increasing ethanol gradient (5min in 30%, 50%, 70%, 95%) and 3 times for 10min in absolute ethanol. Impregnation was performed with Epon A (75%) plus Epon B (25%) plus DMP30 (1,7%). Inclusion was obtained by polymerization at 60°C for 72H. Ultrathin sections (approximately 70 nm thick) were cut on a UC7 (Leica) ultra-microtome, mounted on 200 mesh copper grids coated with 1:1,000 polylysine, and stabilized for 1 day at room temperature (RT) and contrasted with uranyl acetate and lead citrate. Sections were acquired with a Jeol 1400JEM (Tokyo,Japan) transmission electron microscope, 80Kv, equipped with a Orius 600 camera and Digital Micrograph.

### *Drosophila* model

Shot GD9507 UAS-RNAi line from VDRC collection crossed to Mef2-GAL4 driver has been used to attenuate shot gene expression specifically in muscles. Third instar larvae were dissected in physiological salt with 25mM EDTA. Body wall muscles were fixed with 4% formaldehyde in PBS for 15min and then rinsed three times for 5min each in PBS with 0.5% Tween 20 (PBT). Muscles were blocked for 30min with 20% horse serum in PBT at RT. Staining was performed by using primary antibodies applied overnight at 4°C and after washing 3 times in PBT secondary antibodies were applied at RT for 1H. The following primary antibodies were used: anti-Brp1 (1:100; DSHB, Nc82-s), anti-Shot (1:100; DSHB, mAbRod1).

### *Mouse* model

The following mice have been described previously, mice that carry a loxP-flanked allele of *Macf1* (Goryunov et al., 2010) and Hsa-Cre transgenic mice (Miniou et al., 1999). All of the experiments and procedures were conducted in accordance with the guidelines of the local animal ethics committee of the University Claude Bernard – Lyon 1 and in accordance with French and European legislation on animal experimentation and approved by the ethics committee CECCAPP and the French ministry of research. At least 3 *Macf1*^*f/f*^ Cre- or Cre+ mice were tested for each experiment.

### Genotyping of mice

In order to extract the DNA, samples (mice tales) were incubated in extraction solution (25mM NAOH – 0.2mM EDTA) for 30min at 95°C. Following the addition of neutralization solution (40mM Tris-HCL), samples were centrifuged at 13000RPM for 2min at RT and the DNA were collected. DNA concentration was evaluated using Nanodrop (ThermoFisher Scientifics). PCR were carried (Hot Start Taq polymerase (QIAGEN, 1007837), PCR buffer mix (QIAGEN, 1005479) DNTP mix (Biolabs, 447)) using following primers. The same amount of PCR products were then loaded in 2X Agarose-1X TAE gels and were migrated for 2H at 130V. Results were obtained using GelDoc (BioRad).

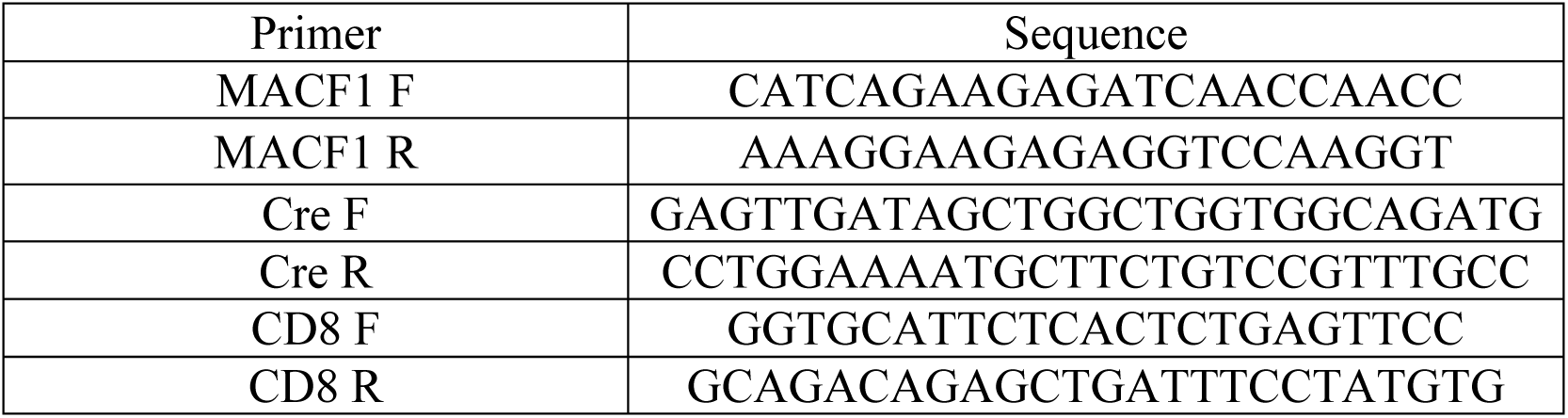

### RNA extraction

After the addition of Trizol (Sigma, T9424-200mL) on each sample, lysing matrix D and fast prep system (MPbio, 6913-100) were used for sample digestion and pre-RNA extraction. In order to extract RNA, samples were incubated in chloroform for 5min at RT, centrifuged for 15min at 12000 rcf at 4°C and incubated in the tubes containing isopropanol (precipitatation of RNA) for 10min at RT. following a centrifuge of samples for 15min at 12000rcf at 4°C, samples were washed 2 times with 70% ethanol and the final RNA pellets were diluted in ultra-pure RNase free water (Invitrogen, 10977-035). RNA concentration was calculated using Nanodrop (ThermoFisher Scientifics).

### RT-q-PCR

Goscript Reverse Transcriptase System (Promega, A5001) was used, as described by the manufacturer to produce the cDNA. Fast Start Universal SYBR Green Master (Rox)(Roche, 04913914001) and CFX Connect™ Real-Time PCR Detection System (BioRad) were used to carry out the quantitative PCR using the following primer sets. The CT of target genes were normalized on 3 control genes.

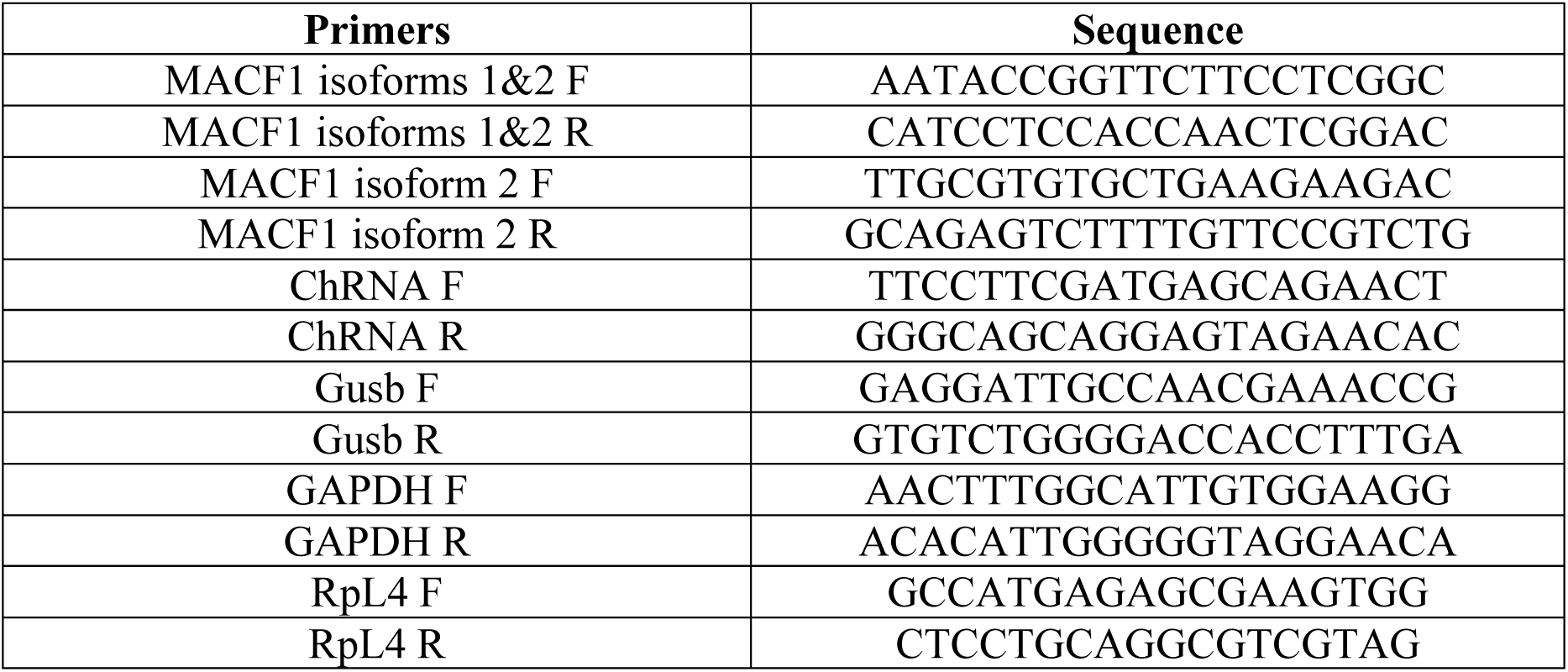

### Histological staining and analysis

*Tibialis anterior, Soleus* and *Gastrocnimius* muscles were collected, embedded in tragacanth gum, and quickly frozen in isopentane cooled in liquid nitrogen. Cross-sections (10µm thick) were obtained from the middle portion of frozen muscles and processed for histological, immunohistochemical, enzymohistological analysis according to standard protocols. The fibre cross-sectional area and the number of centrally nucleated fibers were determined using Laminin and Dapi-stained sections. Fluorescence microscopy and transmission microscopy were performed using Axioimager Z1 microscope with CP Achromat 5x/0.12, 10x/0.3 Ph1, or 20x/0.5 Plan NeoFluar objectives (Zeiss). Images were captured using a charge-coupled device monochrome camera (Coolsnap HQ, Photometrics) or color camera (Coolsnap colour) and MetaMorph software. For all imaging, exposure settings were identical between compared samples. Fiber number and size, central nuclei and peripheral myonuclei were calculated using ImageJ software.

### Succinate DeHydrogenase (SDH) staining and analysis

SDH staining was performed as previously described (NACHLAS et al., 1957). Briefly, transverse sections (8µm) were cut from the mid-belly of the TA muscles on a cryostat at −20°C and stored at −80°C until SDH staining was performed. The sections were dried at room temperature for 30min before incubation in a solution made up of 0.2M phosphate buffer (pH 7.4), 0.1M MgCl2, 0.2M Succinic Acid (Sigma) and 2.4mM NitroBlue Tetrazolium (NBT, Sigma) at 37°C in a humidity chamber for 45min. The sections were then washed in deionized water for 3min, dehydrated in 50% ethanol for 2min, and mounted for viewing with DPX mount medium (Electron Microscopy Sciences). Images were acquired as described above.

### Confocal imaging of T-tubule network

*FDB* muscle fibers were incubated for 30min in the presence of 10µm di-8-anepps in Tyrode solution. Estimation of the T-tubule density from the di-8-anepps fluorescence was carried out from a largest possible region of interest excluding the plasma membrane, within each fiber. For each fiber, two images taken at distinct locations were used. Analysis was carried out with the ImageJ software (National Institute of Health). Automatic threshold with the Otsu method was used to create a binary image of the surface area occupied by T-tubules. The “skeletonize” function was then used to delineate the T-tubule network. T-tubule density was expressed as the percent of positive pixels within the region. Sarcomere length was estimated from half the number of fluorescence peaks (T-tubules) along the length of the main axis of a given fiber. To assess variability in T-tubule orientation, objects within two T-tubule binary images of each fiber were outlined and particle analysis was performed to determine the angle of all objects yielding a perimeter larger than an arbitrary value of 10µm. For each fiber, the standard deviation of angle values was then calculated. This analysis was performed on 10 muscle fibers from 3 Macf1f/f Cre- and from 3 Macf1f/f Cre+ mice, respectively.

### Intracellular Ca^2+^ in voltage-clamped fibers

Single fibers were isolated from *FDB* muscles as described previously (Jacquemond, 1997). In brief, muscles were incubated for 60min at 37 °C in the presence of external Tyrode containing 2 mg.mL collagenase (Sigma, type 1). Single fibers were obtained by triturating the collagenase-treated muscles within the experimental chamber.

Isolated muscle fibers were handled with the silicone voltage-clamp technique (Lefebvre et al., 2014). Briefly, fibers were partly insulated with silicone grease so that only a short portion (50-100µm long) of the fiber extremity remained out of the silicone. Fibers were bathed in a standard voltage-clamp extracellular solution containing (in mM) 140 TEA-methanesulfonate, 2.5 CaCl2, 2 MgCl2, 1 4-aminopyridine, 10 HEPES and 0.002 tetrodotoxin. An RK-400 patch-clamp amplifier (Bio-Logic, Claix) was used in whole-cell configuration in combination with an analog-digital converter (Axon Instruments, Digidata 1440A) controlled by pClamp 9 software (Axon Instruments). Voltage-clamp was performed with a micropipette filled with a solution containing (in mM) 120 K-glutamate, 5 Na2-ATP, 5 Na2-phosphocreatine, 5.5 MgCl2, 15 EGTA, 6 CaCl2, 0.1 rhod-2, 5 glucose, 5 HEPES. The tip of the micropipette was inserted through the silicone within the insulated part of the fiber and was gently crushed against the bottom of the chamber to ease intracellular equilibration and decrease the series resistance. Intracellular equilibration of the solution was allowed for 30min before initiating measurements. Membrane depolarizing steps of 0.5s duration were applied from -80mV. Confocal imaging was conducted with a Zeiss LSM 5 Exciter microscope equipped with a 63x oil immersion objective (numerical aperture 1.4). Rhod-2 fluorescence was detected in line-scan mode (x,t, 1.15 ms per line) above 560nm, upon excitation from the 543nm line of a HeNe laser. Rhod-2 fluorescence transients were expressed as F/F0 where F0 is the baseline fluorescence. The Ca2+ release flux (rate of SR Ca2+ release) was estimated from the time derivative of the total myoplasmic Ca2+ ([Catot]) calculated from the occupancy of intracellular calcium binding sites following a previously described procedure (Kutchukian et al., 2017).

### *In vivo* force measurements

Mice were initially anesthetized in an induction chamber using 4% isoflurane. The right *Hindlimb* was shaved before an electrode cream was applied at the knee and heel regions to optimize electrical stimulation. Each anesthetized mouse was placed supine in a cradle allowing for a strict standardization of the animal positioning. Throughout a typical experiment, anesthesia was maintained by air inhalation through a facemask continuously supplied with 1.5% isoflurane. The cradle also includes an electrical heating blanket in order to maintain the animal at a physiological temperature during anesthesia. Electrical stimuli were delivered through two electrodes located below the knee and the Achille’s tendon. The right foot was positioned and firmly immobilized through a rigid slipper on a pedal of an ergometer (NIMPHEA_Research, AII Biomedical SAS) allowing for the measurement of the force produced by the Hindlimb muscles (i.e., mainly the *Gastrocnemius* muscle). The right knee was also firmly maintained using a rigid fixation in order to optimize isometric force recordings. Monophasic rectangular pulses of 0.2ms were delivered using a constant-current stimulator (Digitimer DS7AH, maximal voltage: 400V). The force-frequency curves were determined by stepwise increasing stimulation frequency, with resting periods > 30s between stimuli in order to avoid effects due to fatigue. For each stimulation train, isometric peak force was calculated. After a 3-min recovery period, force was assessed during a fatigue protocol consisting of 30Hz stimulation trains of 0.3s delivered once every second for 180s. The peak force of each contraction was measured and averaged every 5 contractions. A fatigue index corresponding to the ratio between the last five and the first five contractions was determined. Force signal was sampled at 1000Hz using a Powerlab system and Labchart software (ADinstruments).

### Quantification methods for myonuclei spreading in myotubes

Quantifications in immature myotubes were assessed using an analysis tool developed in our team. An image analysis performed in ImageJ® software is combined with a statistical analysis in RStudio® software. This provides quantifications of parameters, ranked by myonuclei content per myotubes, regarding phenotype of myotubes (area, length) and their respective myonuclei positioning compare to centroid of myotubes (DMcM).

MSG diagrams were obtained through the normalization of lengths of all analyzed myotubes (independently to their myonuclei content) to 100%. White lines represent myonuclei density curves assessing the statistical frequency for myonuclei positioning along myotubes. Each color group reflects statistical estimation of myonuclei clustering along myotubes.

### Quantification methods for EB1 Comets

To determine EB1 comets speed, four continuous frame of a time-lapse movie of EB1-GFP were overlapped, the first two are color-coded in green and the last two are color-coded in red. Comets length was counted if green and red length were equal, traducing a comets growing in the same focal plan.

### Quantification of mitochondria fragmentation

In order to analyze mitochondria repartition, Cytochrome-C staining was used as representative of the mitochondrial network in long differentiated myofibers. Representative images were taken from myofibers and the analysis of images was done using ImageJ® software. One or several Regions Of Interest (ROI) were selected per image, these regions were set just next-to or close to myonuclei. Following a threshold determination, the whole information of each stained particle (representing a single or a group of mitochondria) was extracted from ImageJ software. The particles with circularity equivalent to 1 were eliminated in order to purify our results from any unwanted background errors. Particles with feret less than 0.75µm (mean size of mitochondria) were eliminated to purify our results. Finally, the ratio of number of particles per area of ROI was calculated.

### Quantification of AChRs fragmentation

Bungarotoxin staining was used to analyze formation of acetylcholine receptors clusters in mature myofibers. Representative images were taken from myofibers of each condition as described before and the analysis of images was done using ImageJ® software. Briefly, a threshold of 8µm was set as the minimal size for AChRs clusters. Myofibers with at least one cluster of 8µm or bigger were considered as positive.

### Statistical analysis

The statistical analyses were performed using Student’s t test or the Mann-Whitney test according to samples amount and distribution

